# Robust and accessible in situ sequencing with an open-source automated sequencer

**DOI:** 10.64898/2026.10.18.759961

**Authors:** Aixin Zhang, Zoe Maltzer, Jacob Lapka, Sabrina Cheng, Alan Guthrie, Cliff Slaughterbeck, Angela Y. Fan, Neha Goswami, Mara C.P. Rue, Maryam Majeed, Dennis M. Maharjan, Jeanelle Ariza, Angela P. Ayala, Madeleine Bixby, Cory T. Dickson, Samantha D. Hastings, Alvin Huang, Tulip E. O’Neill, Alana A. Oyama, Caleb Shi, Jack Waters, Yi-Chen Wu, Renae Galluccio, Huiqing Zhan, James A. Bourne, Gregory Horwitz, Justus M. Kebschull, Ian R. Wickersham, Peter Rudebeck, Alex Cahoon, Xiaoyin Chen

## Abstract

Targeted in situ sequencing underlies many spatial transcriptomic, barcoded connectomic, and multiplexed perturbation techniques. Each experiment requires many cycles of chemical reactions and imaging that are time-consuming and error-prone to perform manually, and few commercial platforms can sequence highly diverse RNA barcodes in situ at cellular resolution across large tissue sections. Here we present Seq-o-Matic, an open-source automated in situ sequencer built entirely from commercially available components, which integrates fluidics, heated incubation, and imaging of up to three continuous 18 mm × 50 mm areas on standard microscope slides. Relative to manual sequencing, Seq-o-Matic reduced hands-on time 6-to 17.5-fold and total run time about 2.6-fold. Across three runs spanning two mouse strains and a neonatal marmoset, sequencing quality, measured by signal retention, channel specificity, and phasing, was similar to that of four manually sequenced mouse brains. In four mouse hemispheres sequenced in separate runs (two C57BL/6J, one NOD, and one WSB), consensus cell types were present in every brain and corresponded to those of a manually sequenced C57BL/6J brain. Within cortical excitatory neurons, the two C57BL/6J brains were more similar to each other than were brains of different strains, suggesting that technical variation between runs was small relative to strain differences. Seq-o-Matic thus makes consistent, high-throughput in situ sequencing accessible to laboratories without specialized in situ sequencing expertise.

## Introduction

In situ sequencing reads out RNA sequences directly in tissue at submicron resolution, and its targeted form (Ke et al., 2013) has become the foundation of a growing family of techniques. Targeted in situ sequencing can operate in two modes: In the first mode, a probe binds directly to its target transcript and reports its presence, which allows simultaneous detection of endogenous transcripts from hundreds of genes or more (Alon et al., 2021; Bugeon et al., 2022; X. Chen et al., 2025; Wang et al., 2018), much like highly multiplexed in situ hybridization techniques (Borm et al., 2023; K. H. Chen et al., 2015; Codeluppi et al., 2018; Eng et al., 2019). In the second mode, the target sequence is copied onto the probe itself. This allows the method to read out highly diverse synthetic RNA barcodes de novo, which hybridization-based techniques cannot do because they detect only predefined sequences. This capability underlies barcoded connectomics (X. Chen et al., 2019; Y. Chen et al., 2022; Majeed et al., 2026; Sun et al., 2021; Yuan et al., 2024; Zhang et al., 2024), in which diverse barcodes uniquely label individual neurons. Because these techniques all build on in situ sequencing, making in situ sequencing itself accessible would extend all of them to laboratories with diverse expertise and questions.

In principle, in situ sequencing can be set up in any laboratory, because it relies on commercially available reagents and equipment (X. Chen et al., 2018; Feldman et al., 2019). In practice, however, it is labor-intensive and error-prone. After RNAs in a tissue section are converted into amplified sequencing templates (library preparation), the sample undergoes many cycles of sequencing reactions and imaging; each cycle sequentially reads out one position from the RNAs being sequenced. To resolve a panel of hundreds of genes, a typical experiment requires seven sequencing cycles; when RNA barcodes are also sequenced, as in barcoded connectomics techniques, more than 20 cycles are required (**<u>Table S1</u>**).

Because each cycle builds on the previous one, an error in any cycle compromises all subsequent cycles. Automating in situ sequencing would address these problems, but existing commercial platforms do not meet the needs of barcoded connectomics approaches. Capture-based platforms, such as Stereo-seq, 10x Visium, and Illumina StrataMap, and their open-source counterparts, such as Seq-Scope and Open-ST (Cho et al., 2021; Schott et al., 2024), capture molecules only at the tissue surface. Because some barcoded connectomics techniques require reading out RNAs from axons (Majeed et al., 2026; Yuan et al., 2024) or from all cell bodies distributed throughout the section (Zhang et al., 2024), surface capture is insufficient. Imaging-based platforms, such as 10x Xenium and Atera, detect transcripts with predefined probes and therefore cannot read out sequences de novo. The Singular Genomics G4X can read sequences de novo, but it accommodates only small samples and precludes intact sections from large brains.

Moreover, even a commercial system that met all of these requirements would still require a substantial upfront investment in a dedicated instrument. An automated in situ sequencer that reads sequences de novo across large, intact sections, and that can be built at modest cost around an existing motorized microscope, would therefore lower the barrier to entry for BARseq and other in situ sequencing techniques, including for laboratories that cannot justify a dedicated instrument.

Here we present Seq-o-Matic, an open-source automated in situ sequencer that meets these requirements. Seq-o-Matic performs all sequencing reactions and imaging automatically on standard microscope slides, substantially reducing hands-on and total time. Because Seq-o-Matic controls the microscope through open-source software, laboratories can add it to an existing compatible motorized microscope. In addition, the imaging area on each slide is large enough to accommodate multiple intact hemi-coronal sections from a neonatal marmoset brain. Across mouse and marmoset tissue, Seq-o-Matic achieved sequencing quality similar to that of manual sequencing, and data from separately sequenced brains were consistent enough to resolve differences between mouse strains. Because Seq-o-Matic is built entirely from commercially available components and open-source software, it removes the engineering and programming expertise otherwise needed to build an automated sequencer, lowering a major barrier to in situ sequencing for non-expert laboratories.

## Results

### Seq-o-Matic automates in situ sequencing with open-source hardware and software

To reduce labor and error-prone steps from in situ sequencing, we designed and built *Seq-o-Matic*, an automated in situ sequencer that integrates all sequencing reactions and imaging steps. For a typical experiment, Seq-o-Matic reduces days of pipetting and imaging to a two-hour initial setup. Compared with manual sequencing, automation reduces total run time 2.6-to 2.7-fold and hands-on time 6-to 17.5-fold, depending on the number of cycles (**<u>Table S1</u>**).

Seq-o-Matic integrates fluidic control, imaging, and data preprocessing (**<u>Fig. 1A</u>**). The user provides a protocol file that specifies the sequence of cycles to be performed, and Seq-o-Matic executes each step through two modules that separately control fluidics and imaging. For reaction steps, the fluidics module performs fluid exchange and heated incubations as specified in recipe files. For imaging steps, the imaging module, which controls the microscope through Micro-Manager (Edelstein et al., 2014) via its Python interface Pycro-Manager (Pinkard et al., 2021), refocuses, corrects slide positions for any drift relative to the initial positions, and images all tiles covering the regions of interest. As each tile is acquired, it is converted to a maximum-intensity projection, which is stored locally and uploaded to a server or cloud storage. In our system, Seq-o-Matic drives a spinning disk confocal microscope with a large field of view, an objective piezo focus for fast focusing, and a long-working distance objective (see below), but in principle, Seq-o-Matic can be used with any fully motorized microscope that Micro-Manager supports. To make experiments reproducible, imaging conditions (laser power, exposure, and camera settings) and fluidic recipes cannot be edited from the graphical user interface; instead, they are stored in JSON files that advanced users can modify for custom experiments.

**Figure 1.**
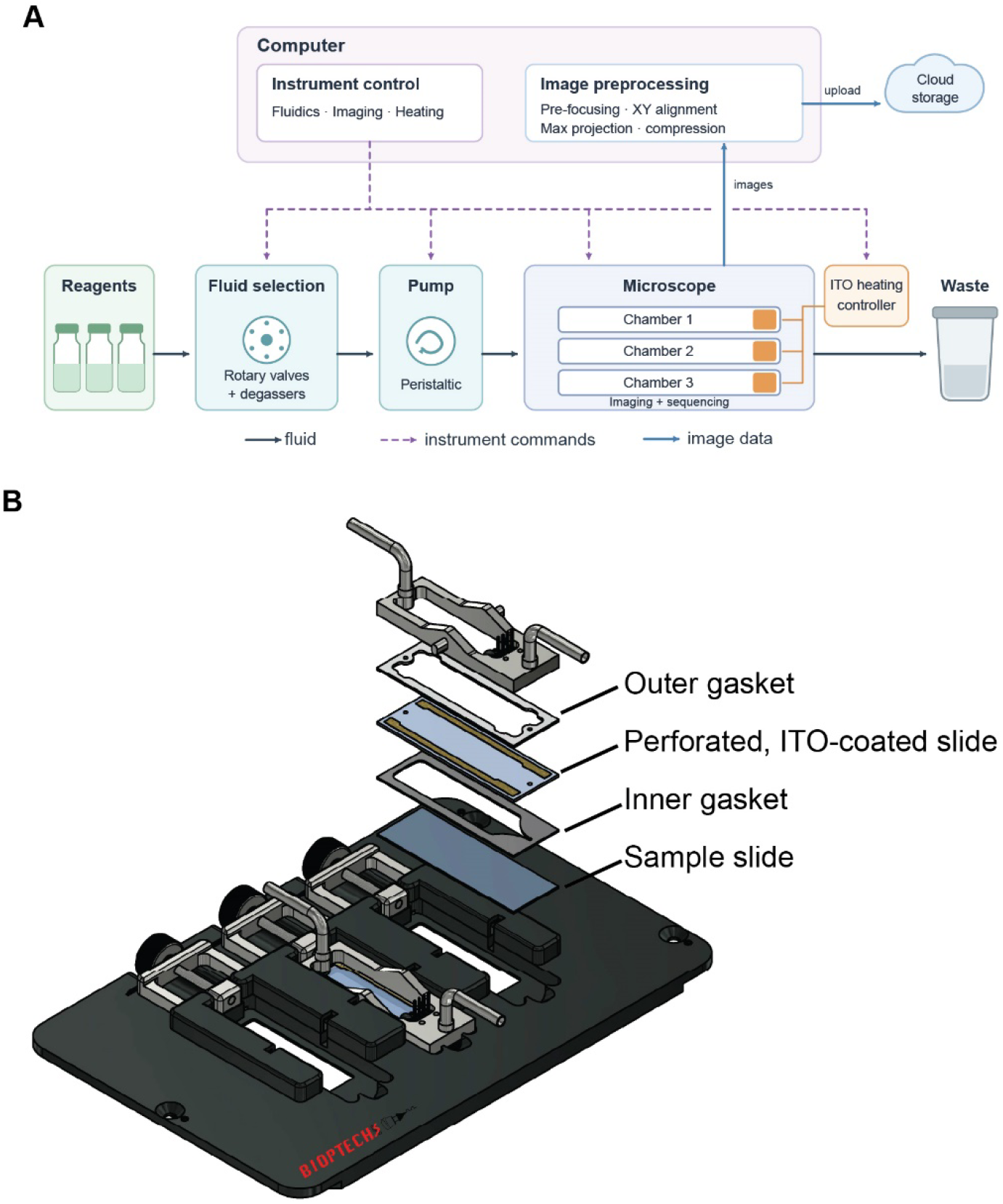
Seq-o-Matic automates in situ sequencing on standard slides. **(A)** Schematic of Seq-o-Matic. The control computer sends instrument commands (dashed purple arrows) to the fluid selection module (rotary valves and degassers), the pump, the microscope, and the ITO heating controller. Reagents (black arrows) are pumped through three sample chambers on the microscope stage and then to waste. Images (blue arrows) are pre-focused, aligned in XY, maximum-projected, and compressed on the computer and uploaded to network storage. **(B)** Exploded view of one sample chamber in the three-slide stage insert. Each chamber consists of, from bottom to top, the sample slide, an inner gasket, a perforated ITO-coated slide, an outer gasket, and the chamber top with inflow and outflow tubes.

Samples are held in a commercial heated perfusion chamber (Bioptechs Tri-Imaging Chamber) that is optimized for in situ sequencing. This chamber accommodates up to three standard 25 mm × 75 mm × 1 mm microscope slides, each with an 18 mm × 50 mm imaging window (**<u>Fig. 1B</u>**). Each window can hold multiple mouse hemi-coronal sections or intact hemi-coronal sections of neonatal marmoset brain.

Mounting tissue on standard slides rather than coverslips reduces the risk of sample loss from breakage. This is particularly important for axonal BARseq (Majeed et al., 2026; Yuan et al., 2024) and barcoded rabies tracing (Becalick et al., 2026; Zhang et al., 2024), in which every section from a tissue block may need to be sequenced. In each chamber, a silicone gasket separates the sample slide from a perforated slide coated with indium tin oxide (ITO). The transparent ITO layer permits transmitted-light illumination and heats the chamber resistively when current is applied. Samples are imaged through the 1-mm-thick slides with a long-working-distance objective that corrects for slide thickness (Nikon CFI S Plan Fluor LWD 20XC, 0.70 NA). The three-slide assembly fits the footprint of a standard K-frame stage insert, so it can be installed on a broad range of microscope stages with appropriate adapters.

All components are commercially available (**Table S2**), and we provide the complete software package (https://github.com/BarseqLab/Seq_o_matic.git) along with a detailed hardware installation and operation guide (see **Supp File 1**, also at https://github.com/BarseqLab/Seq_o_matic/blob/main/docs/setup_guide/Seq-o-matic_Setup_Guide.md).

### Seq-o-Matic achieves high sequencing quality

To assess sequencing quality, we sequenced one hemisphere each from an adult male NOD/ShiLtJ mouse (37 hemi-coronal sections) and an adult male WSB/EiJ mouse (34 hemi-coronal sections), as well as 22 hemi-coronal sections from a postnatal day 0 (PD0) common marmoset. Because each imaging window holds multiple sections (**<u>Fig. 2A, B</u>**), each dataset fit on three slides and was sequenced in a single run in under two days. We compared these data with four P28 C57BL/6J mouse brains that we previously sequenced manually on the same microscope (X. Chen et al., 2025). We did not match donors in this comparison, because raw sequencing performance is expected to depend primarily on the sequencing chemistry and the instrument, not donor type. The mouse samples were sequenced with a 104-gene cell-type marker panel (X. Chen et al., 2025) and the marmoset sample with a 60-gene panel (**<u>Table S3</u>**); both panels were designed to distinguish cortical excitatory neuron types (X. Chen et al., 2025; Fan et al., 2026). Together, these datasets span two species, two mouse strains, and two gene panels, allowing us to test whether Seq-o-Matic maintains consistent sequencing performance across diverse samples.

**Figure 2.**
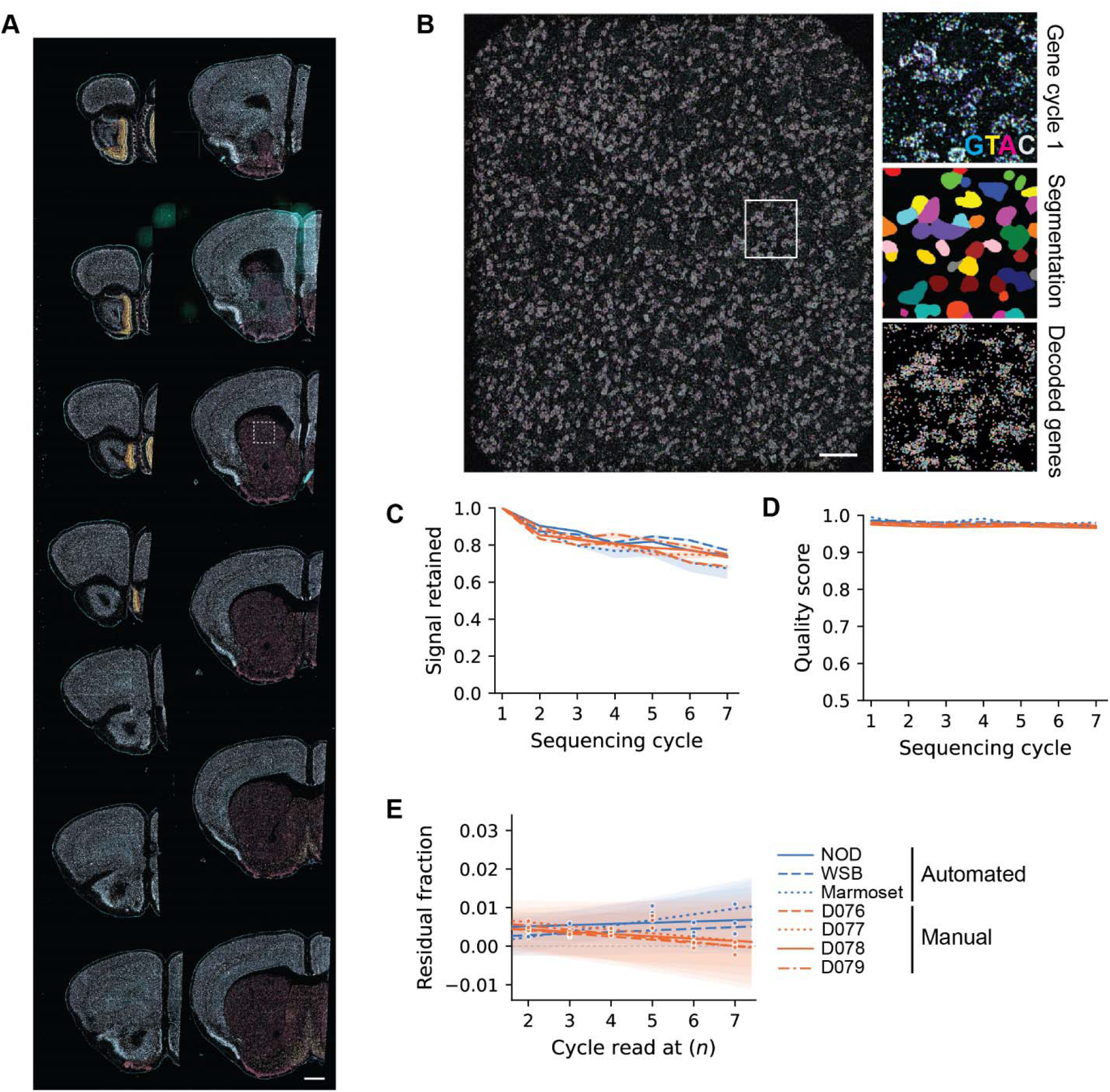
Seq-o-Matic achieves sequencing quality comparable to manual sequencing. **(A)** A representative slide showing 14 hemi-coronal sections from the NOD brain during the hybridization cycle. Cyan = *Slc17a7*, Yellow = *Gad1*, White = *Slc30a3*, and Magenta = all other 101 genes that were decoded through sequencing. The dashed box marks the region shown in B. Scale bar, 1 mm. **(B)** *Left*, one image tile from the boxed region in A. *Right*, the boxed region on the image on the left at higher magnification, showing the first gene sequencing cycle (G, T, A, and C channels), cell segmentation, and decoded gene rolonies. Scale bar, 100 µm. **(C)** Fraction of the first-cycle signal retained over seven sequencing cycles. For each rolony signal, the intensity of the called base in each cycle was divided by its first-cycle intensity; values are medians across rolony signals within each image tile and then across tiles. Each line is one brain: blue, sequenced with Seq-o-Matic (NOD, WSB, and marmoset); orange, sequenced manually (four C57BL/6J brains) (X. Chen et al., 2025). Line styles identify brains as in (E). Shading, 95% bootstrap confidence intervals over image tiles. **(D)** Quality score (intensity of the brightest channel divided by the square root of the summed squared intensities of all four channels; 0.5, equal signal in all channels; 1, signal in one channel) in each sequencing cycle, shown as in (C). **(E)** Phasing.

A key measure of raw sequencing quality is how well rolony (rolling-circle amplification product) fluorescence is preserved across sequencing cycles. Signal can decline progressively through loss of template, DNA damage, and accumulated chemical scarring that leaves sequencing reactions incomplete. To quantify this decline, we identified all rolony signals (local intensity maxima) in the first cycle, whether or not they were later decoded, and tracked the intensity of the called base at each position through seven cycles. Because the called base can change channel between cycles, we first normalized each channel to its median first-cycle intensity and applied the same scaling to that channel in all cycles. For each rolony signal, we then divided the called-base intensity in each cycle by its first-cycle intensity, and summarized retention as the median across signals within each image tile and then the median across tiles. At cycle 7, the three brains sequenced with Seq-o-Matic retained 67.4% to 77.2% of their first-cycle signal, compared with 68.5% to 75.0% for the four manually sequenced brains (X. Chen et al., 2025) (**<u>Fig. 2C</u>**). Signal retention with Seq-o-Matic was thus similar to that of manual sequencing.

Because each nucleotide incorporated in Illumina MiSeq chemistry carries a single fluorophore, an ideal reaction produces signal in only one of the four fluorescence channels for each rolony. We quantified this channel specificity for each decoded rolony using a standard in situ sequencing quality score: the intensity of the brightest channel divided by the square root of the summed squared intensities of all four channels (X. Chen et al., 2018, 2019; Lee et al., 2014). The score ranges from 0.5, when all four channels are equally bright, to 1, when all signal is in one channel. Throughout seven cycles, the median score remained at or above 0.97 in the three brains sequenced with Seq-o-Matic and in the four manually sequenced brains (X. Chen et al., 2025), with values of 0.97 to 0.98 at cycle 7 in both groups (**<u>Fig. 2D</u>**). Automation thus did not detectably reduce channel specificity.

Because each rolony contains many copies of the same template, some molecules may fail to extend during a cycle and thereafter lag one base behind. This phasing accumulates across cycles. If *p* is the per-cycle probability that a molecule fails to extend, then after *n* cycles the fraction of molecules lagging by exactly one base is *np*(1 − *p*)^(*n* − 1), and the in-phase fraction in the preceding cycle is (1 − *p*)^(*n* − 1). Their ratio, *R*(*n*) = *np*, therefore increases linearly with *n*, and its slope estimates *p*. To estimate *R*(*n*), we analyzed decoded rolonies whose base call changed between consecutive cycles and measured the residual signal in the previous base’s channel, relative to that channel’s intensity in the preceding cycle, after subtracting a brightness-matched background (**<u>Methods</u>**). To avoid confounding lagging signal with channel bleed-through, we excluded the four base changes within bleed-through pairs (G→T, T→G, A→C, and C→A) and used the remaining eight. We fitted *R*(*n*) against *n* by linear regression and estimated 95% confidence intervals with a hierarchical bootstrap over image tiles and base changes (2,000 iterations). The phasing slope was below 0.15% per cycle in all seven brains, and the confidence interval included zero in all but the marmoset dataset, which had the steepest slope (0.148% per cycle; 95% confidence interval, 0.034% to 0.291%; **<u>Fig. 2E</u>**). Even in the marmoset, therefore, only about 1% (95% confidence interval, 0.2% to 2%) of the signal lagged by one base after seven cycles, indicating nearly complete sequencing reactions. Assuming a constant per-cycle rate, this would correspond to about 2% lagging signal (95% confidence interval, 0.5% to 4.4%) after the 15 cycles typically used for barcode sequencing (**<u>Table S1</u>**). Together with the similar signal retention and channel specificity, these results indicate that Seq-o-Matic achieves sequencing quality similar to that of manual sequencing.

Residual fraction (signal remaining in the channel of the previous base, relative to that channel’s intensity in the preceding cycle) plotted against the cycle n at which it was read. Dots, median residual fraction across image tiles for each brain and cycle; lines, linear fits; shading, 95% hierarchical bootstrap confidence intervals (2,000 iterations). The slope estimates the per-cycle probability that a molecule fails to extend.

### Seq-o-Matic resolves transcriptomic domains in intact marmoset cortical sections

Seq-o-Matic’s large imaging window allows intact marmoset hemi-coronal sections to be sequenced without physically subdividing the tissue and reconstructing it afterward. We used this capability to ask whether spatial gene-expression patterns could delineate cortical regions, as we previously showed in mouse cortex (X. Chen et al., 2025). We analyzed a representative section (**<u>Fig. 3A</u>**), which we matched to AP +4.30 mm in the adult Paxinos marmoset atlas (Paxinos, 2012) using anatomical landmarks, including the shape of the lateral geniculate nucleus and the positions of the superior temporal and occipitotemporal sulci. We partitioned the cortical ribbon into 70 contiguous bins, each spanning the cortical thickness (**<u>Fig. 3B</u>**), summed the gene counts in each bin, smoothed them over neighboring bins, and reduced the Hellinger-transformed expression profiles to 10 principal components. We then used one-dimensional dynamic programming to partition the ordered bins into contiguous transcriptomic domains, and selected the five-domain solution, beyond which additional domains yielded diminishing gains over a resampling null (**<u>Fig. 3C</u>**; **<u>Methods</u>**). From the medial end of the cortex, the domains corresponded to retrosplenial/posterior cingulate cortex, followed by cingulo-parietal cortex (area 31/PE/AIP), MT and surrounding areas (MST/MTC/FST), temporal areas (TE3/TE2/TF), and finally the entorhinal cortex. Transcriptomic segmentation of an intact marmoset section thus yielded broad domains consistent with major atlas-defined cortical regions.

**Figure 3.**
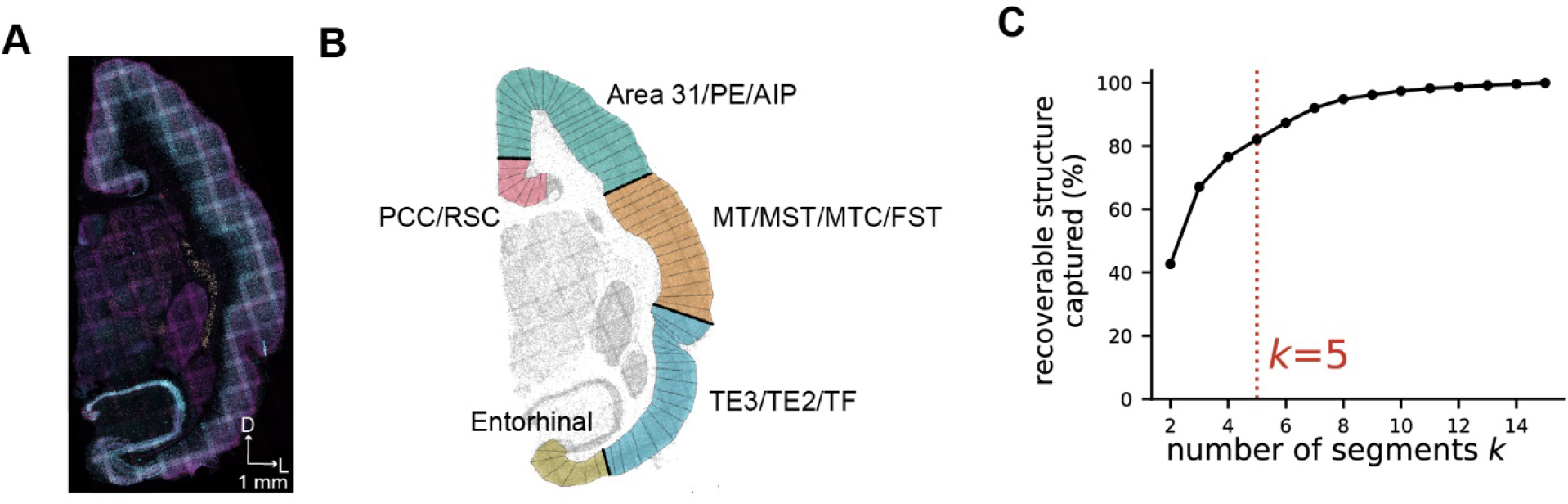
Transcriptomic domains in an intact marmoset hemi-coronal section. **(A)** Hybridization cycle image of a hemi-coronal section from a PD0 marmoset. D, dorsal; L, lateral. Scale bar, 1 mm. **(B)** The cortical ribbon of the section in A, divided into 70 bins that each span the cortical thickness (thin lines). Bins are colored by the five transcriptomic domains identified by one-dimensional dynamic-programming segmentation, with domain borders shown as thick lines. Labels give the atlas regions that correspond to each domain; slashes separate similar areas grouped within a domain. Gray dots, cells. **(C)** Percentage of the recoverable structure captured as a function of the number of domains k. Recoverable structure is the cumulative reduction in within-domain variance beyond that expected by chance, relative to the total up to 15 domains (Methods). Dotted line, the selected five-domain solution.

### Seq-o-Matic data resolve transcriptomic differences across mouse strains

Large-scale studies often combine data from many separately sequenced brains, which requires that technical variation between runs be small relative to the biological differences of interest. As a proof of principle, we tested whether Seq-o-Matic data could resolve transcriptomic differences between mouse strains beyond run-to-run technical variation. In addition to the NOD and WSB brains, we sequenced two C57BL/6J brains with the same gene panel in separate runs and compared neurons across the three strains (**<u>Fig. 4A</u>**). These two brains were sequenced on a second Seq-o-Matic system. Because raw fluorescence intensities and channel bleed-through depend on the imaging hardware, we excluded them from the raw signal analysis above; however, cell-level data, which depend on decoded rolony counts rather than raw intensities, are expected to be comparable across systems.

**Figure 4.**
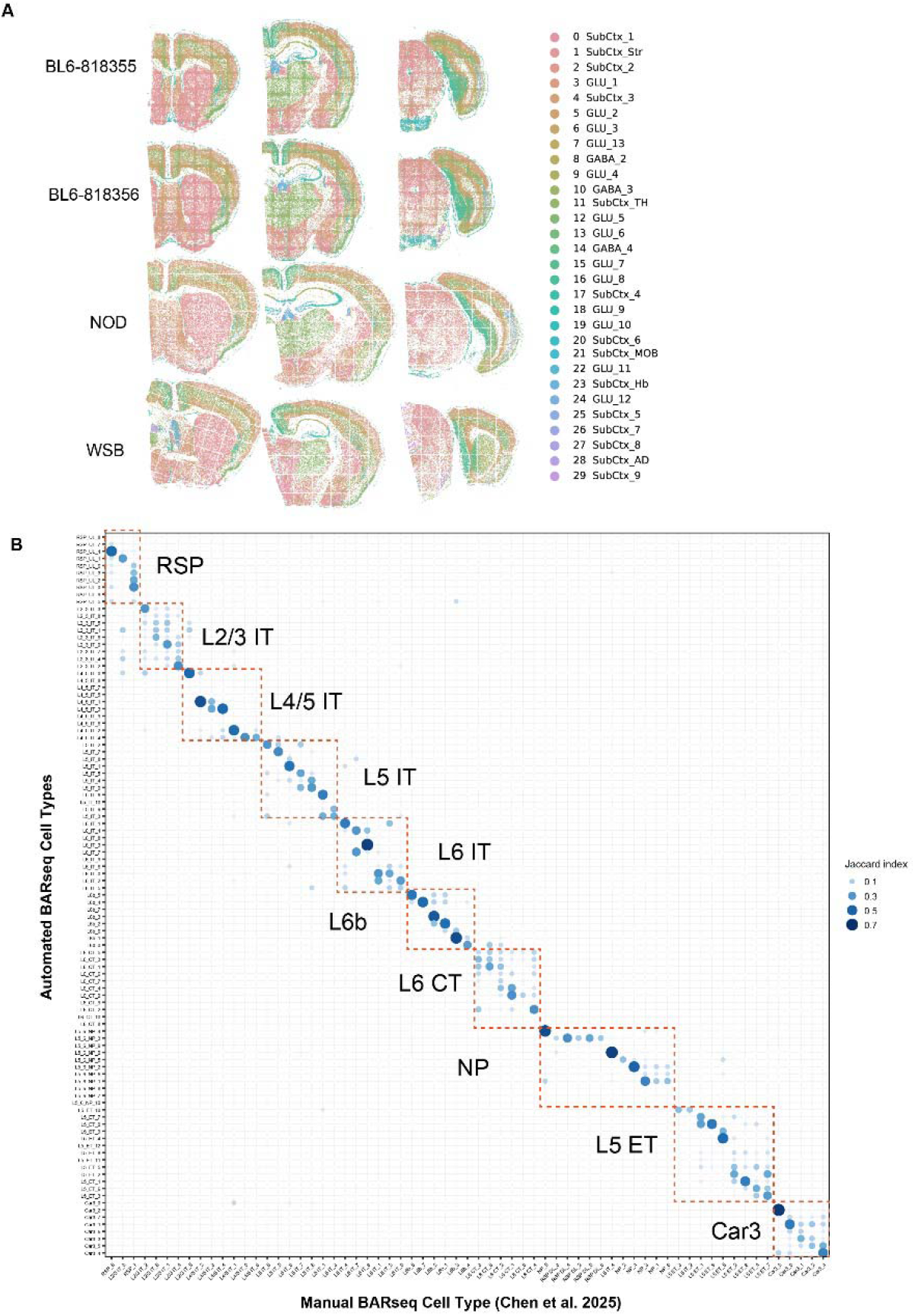
Consensus cell types across four mouse brains sequenced with Seq-o-Matic. **(A)** Three coronal sections from each of the four mouse brains sequenced with Seq-o-Matic (rows: two C57BL/6J, one NOD, and one WSB), with cells colored at the H1 level. **(B)** Overlap (Jaccard index) between the H3 types of cortical excitatory neurons identified in the Seq-o-Matic sequenced C57BL/6J brains (rows) and the H3 types transferred by kNN mapping from a manually sequenced C57BL/6J brain (columns) (X. Chen et al., 2025). Dot size and color indicate the Jaccard index; dashed boxes group H3 types by H2 type.

We first clustered cells hierarchically at three levels to obtain consensus cell-type labels across the four brains (5,619,357 cells with at least 20 counts and 5 genes each), following our previous scheme (X. Chen et al., 2025). We grouped 31 first-level (H1) clusters into excitatory neurons, inhibitory neurons, and other cells, and then re-clustered the excitatory neurons into 16 second-level (H2) types: 10 isocortical and 6 from the hippocampal formation. These H2 types largely corresponded to subclasses in the Allen Brain Cell (ABC) atlas (Yao et al., 2023). Re-clustering each H2 type separately yielded 168 third-level (H3) types, 101 of them cortical. We used Harmony (Korsunsky et al., 2019) to integrate the brains during clustering, so that corresponding cell types received the same labels in every brain. Because Harmony aligns brains by removing differences between them, which are the very differences we aimed to measure, all subsequent analyses used uncorrected data.

To test whether these types matched those identified previously, we transferred H3 labels by k-nearest-neighbor (kNN) mapping from a C57BL/6J brain that we had sequenced manually (465,872 cortical excitatory neurons) (X. Chen et al., 2025) to the excitatory neurons of the two C57BL/6J brains sequenced with Seq-o-Matic, and compared the de novo and transferred labels (**<u>Fig. 4B</u>**). For 89 of 101 cortical H3 types, the best-matching manual type belonged to the same H2 type, and most H3 types matched predominantly to a small number of clusters (**<u>Fig. 4B</u>**); residual overlap with neighboring types is expected, because transcriptomic types within a subclass vary along continuous gradients.

We next used the 10 cortical H2 types to assess reproducibility, with strain differences as a benchmark. Each H2 type was present in all four brains at similar proportions (**<u>Fig. 5A</u>**) and occupied similar laminar positions (**<u>Fig. 5B</u>**), consistent with each H2 type labeling the same cell population in every strain. Within each H2 type, UMAP embeddings showed that neurons from the two C57BL/6J brains largely overlapped, whereas NOD and WSB neurons formed distributions of similar shape but were slightly shifted from the C57BL/6J neurons (**<u>Fig. 5C</u>**). Because the C57BL/6J brains were sequenced on a different system, these shifts could reflect strain differences, run-to-run variation, or differences between systems. We therefore compared two pairs of brains in which both brains were sequenced on the same system, in separate runs: the two C57BL/6J brains, which differ only by run and individual animal, and the NOD and WSB brains, which additionally differ by strain. If run-to-run variation were as large as strain differences, the two C57BL/6J brains should be as distinguishable from each other as the NOD and WSB brains are.

**Figure 5.**
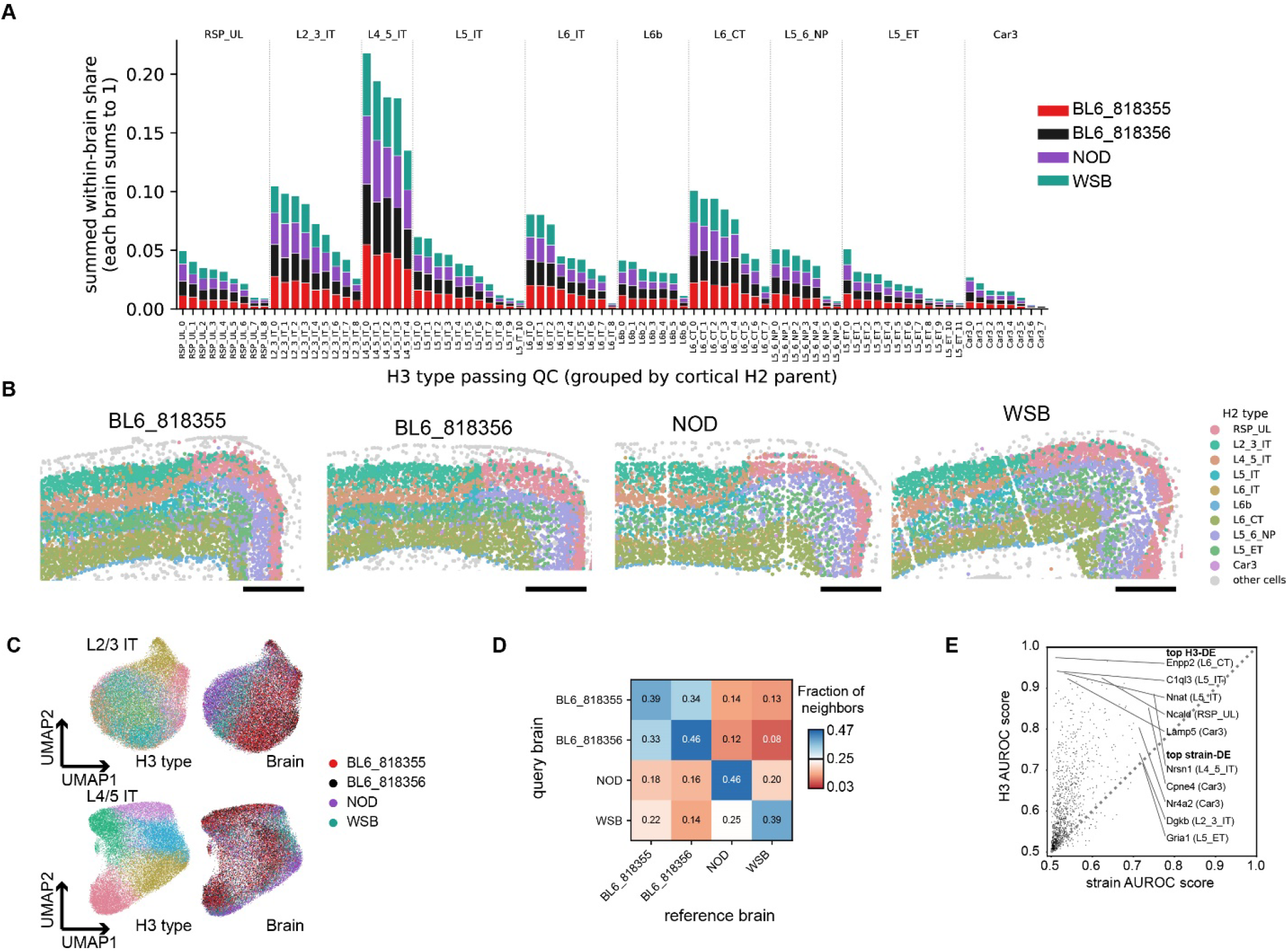
Seq-o-Matic data resolve strain differences in corresponding cortical neuron types. **(A)** Within-brain share of each quality-controlled cortical H3 type, grouped by H2 type and stacked by brain (colors). For each brain, shares sum to 1 across the plotted H3 types. **(B)** Dorsal cortex of one coronal section from each brain, with neurons colored by cortical H2 type and other cells in gray. Scale bars, 500 µm. **(C)** UMAP embeddings, computed without Harmony correction, of neurons of two example H2 types (L2/3 IT and L4/5 IT), colored by H3 type (left of each pair) or by brain (right). **(D)** Fraction of the 40 nearest neighbors of neurons from each query brain (rows) that came from each brain (columns), averaged across the 10 cortical H2 types. Within each H2 type, equal numbers of neurons from the four brains, balanced for H3 type, were pooled, and neighbors were found among all other pooled neurons using the first 20 principal components of log-normalized expression. Colors are centered on the chance level of 0.25 (white); blue, more neighbors than expected; red, fewer. **(E)** AUROC score for distinguishing H3 types within strains (y axis) against that for distinguishing strains within H3 types (x axis), for each gene in each cortical H2 type (one dot per gene and H2 type). Both AUROCs were computed on log-normalized expression, as in (D). AUROCs below 0.5 were reflected about 0.5. Labels mark the five points with the highest H3 score and the five with the highest strain score; dotted line, identity.

To compare them, we used a k-nearest-neighbor analysis (**<u>Fig. 5D</u>**). Within each H2 type, we pooled equal numbers of neurons from the four brains, balanced for H3 type, and found the 40 nearest neighbors of each neuron in the space of the first 20 principal components of log-normalized expression (Methods). With equal contributions, the expected fraction of neighbors from each brain was 0.25. Each brain had the most neighbors within itself, but neurons from each C57BL/6J brain had nearly as many neighbors in the other C57BL/6J brain (same brain versus other C57BL/6J brain: 0.39 versus 0.34 and 0.46 versus 0.33). No other pair of brains exceeded chance. Accordingly, the two C57BL/6J brains mixed more with each other than the NOD and WSB brains did, in all 10 H2 types (mixing index, 0.79 versus 0.53, where 1 indicates interchangeable brains and 0 complete separation; one-sided sign test across H2 types, p = 9.8 × 10^-4^; **<u>Methods</u>**). These results suggest that technical variation between separately sequenced brains was small compared with differences between strains.

We next asked how large the strain differences observable with our gene panel were. Because the H3 types approach the resolution limit at which our data can distinguish clusters of neurons, they provide a natural reference scale: we compared the strain differences captured by the panel with the differences among H3 types captured by the same genes. All 85 cortical H3 types that passed quality control (of 101; Methods) were found in all four brains (**<u>Fig. 5A</u>**). For each gene within each H2 type, we then quantified how well its expression distinguished H3 types within each strain, and how well it distinguished strains within each H3 type, using the area under the receiver operating characteristic curve (AUROC; each H3 type or strain versus the rest; **<u>Fig. 5E</u>**). Many genes distinguished H3 types, with the most informative gene in each H2 type reaching an AUROC of 0.84 to 0.97. Many genes also distinguished strains, but generally less strongly: the most strain-informative gene in each H2 type reached an AUROC of 0.64 to 0.75, and 90% of gene and H2 type combinations were more informative of H3 types than of strains. For example, *Enpp2* distinguished H3 types within L6 CT neurons with an AUROC of 0.97 but distinguished strains with an AUROC of only 0.51. Two features of this comparison favor H3 types: the panel was designed to distinguish cell types, and the H3 types were defined using the same genes. Conversely, because the strain comparison also includes any variation between runs and systems, it may overestimate true strain differences. Within the resolution of our panel, strain differences were thus generally smaller than the differences among fine-grained cell types, yet the neighbor analysis resolved them (Fig. 5D), indicating that Seq-o-Matic data can detect modest transcriptomic differences across separately sequenced brains.

## Discussion

Here we presented Seq-o-Matic, an automated in situ sequencer that achieves data quality comparable to manual sequencing, with low variation across experimental batches. Because Seq-o-Matic is built from commercially available parts around an existing motorized microscope, and its hardware design, software, and protocols are openly available, it makes in situ sequencing and barcoded connectomics accessible at low cost to laboratories with diverse expertise. Moreover, its large imaging window and consistency across experiments open new possibilities for in situ sequencing, from barcoded connectomics in large brains to brain-wide comparisons across strains.

### Automation enables large-scale and large-format in situ sequencing

Automation converts the hands-on labor of in situ sequencing from a cost that scales with the number of sequencing cycles into a largely fixed cost per batch. It also reduces human errors, which can cause sample loss. Moreover, because an error in one cycle affects all subsequent cycles, a single human error can compromise an entire run. The reduced time and reduced errors are particularly important for techniques that require sequencing many sections from a block of tissue with minimal loss. For example, axonal BARseq (Majeed et al., 2026; Yuan et al., 2024) maps the projections of barcoded neurons by matching the barcodes in their somata to those in their axons across the brain, all sequenced in situ; because each lost section removes axons that could otherwise be matched, section loss reduces the sensitivity of projection detection. Similarly, barcoded rabies tracing (Becalick et al., 2026; Zhang et al., 2024) requires sequencing all source cells, because subsampling them introduces ambiguity in assigning presynaptic cells to their postsynaptic partners.

Automation is also key to realizing a central advantage of barcoded connectomics in large brains, such as those of non-human primates. Conventional single-neuron tracing reconstructs each axon by following it continuously through the brain (Peng et al., 2021; Winnubst et al., 2019), so tracing errors accumulate with the distance traced. In contrast, barcoded approaches such as BARseq assign axons to their parent neurons by matching barcodes, and the accuracy of this matching depends on barcode diversity and sequencing accuracy rather than on the distance between soma and axon. Barcoded connectomics is therefore well suited in principle to mapping large brains, but realizing this advantage requires sequencing many large sections routinely. Seq-o-Matic provides both: automation makes the many cycles required for barcode sequencing practical at scale, and its large imaging window accommodates intact hemi-coronal sections of the neonatal marmoset brain on a single slide, so tissue need not be subdivided and computationally reassembled. Together, these features provide a path to barcoded connectomics in larger brains.

### An open-source platform for democratized in situ sequencing

Seq-o-Matic was designed to be built and used by laboratories without prior expertise in in situ sequencing. Seq-o-Matic controls the microscope through Micro-Manager, and can therefore be adapted to other motorized microscopes that are compatible. Although we designed Seq-o-Matic for scalable BARseq experiments, the sequencing chemistry, imaging settings, and experimental design can be customized for each application. In a BARseq experiment, reagents cost about $500 to $1,000 per slide, substantially less than per-slide charges for most commercial platforms. Beyond cost, Seq-o-Matic allows laboratories to pilot in situ sequencing on a microscope they already own before investing in a dedicated instrument. To date, at least five Seq-o-Matic systems have been built from the components and documentation provided here: three at the Allen Institute, including the two that produced the data in this study; one at the MAPseq Core Facility at Cold Spring Harbor Laboratory; and one at Johns Hopkins University. The reproduction of the instrument at three institutions suggests that it can be built and operated beyond the laboratory in which it was developed. By making the hardware design, software, and protocols openly available, we thus aim to democratize barcoded connectomics and in situ sequencing, so that laboratories with diverse expertise can apply these techniques to their own questions.

### Low run-to-run variation allows comparisons across strains

Comparing samples across experiments requires that technical variation between experiments not obscure the biological differences of interest. Here, Seq-o-Matic data resolved transcriptomic differences between mouse strains in corresponding cortical neuron types, even though these differences were generally smaller than those among fine-grained cell types. This comparison was a stringent test, because our gene panel consisted of marker genes chosen to distinguish cortical cell types (X. Chen et al., 2025), not genes that differ across strains; the panel therefore carried less strain signal than a strain-informed one would. In addition, two separately sequenced C57BL/6J brains were nearly interchangeable in a neighbor analysis, whereas brains from different strains were not. These results suggest that Seq-o-Matic data can be compared across experiments to reveal strain-to-strain differences on a brain-wide scale.

Because the gene panel was not optimized to capture strain-to-strain differences, the differences we observed should not be read as the full extent of transcriptomic variation across these strains. Adding genes that are differentially expressed across strains, identified for example by single-cell RNA sequencing of the same strains, would allow more sensitive and comprehensive detection of strain differences. Since in situ sequencing preserves the positions of neurons, the same approach also provides a path to examine anatomical differences across strains, such as differences in laminar positions, spatial distributions, and areal organization of cell types.

## Methods

### Animals

#### Mouse

Animal handling was conducted according to protocols approved by the Institutional Animal Care and Use Committee (IACUC) of the Allen Institute. Animals were housed 3-5 per cage and were on a 12/12 light/dark cycle in an environmentally controlled room (humidity 40%, temperature 21 °C). 8 to 12 week old male C57BL/6J, NOD/ShiLtJ, and WSB/EiJ animals were purchased from Jackson Laboratory.

#### Marmoset

All experimental procedures were fully compliant with the Guidelines for the Care and Use of Laboratory Animals by the National Institutes of Health and approved by the Animal Care and Use Committee of the National Institute of Mental Health. Animals were housed at ambient conditions (26 - 28 °C, 40 – 60 % humidity) on a circadian cycle of 12-h light and 12-h dark, ad-libitum access to food and water.

### Tissue preparation

Animals were deeply anesthetized with isoflurane and transcardially perfused through the left ventricle with 0.9% saline. Brains were dissected and embedded in OCT, then frozen in an ethanol-dry ice bath.

The brains were then stored at −80 °C, then cryosectioned to 20 µm and mounted onto Nexterion A+ slides. The slides were stored at −80 °C until library preparation.

### In situ library preparation

BARseq library preparation was performed as previously described (X. Chen et al., 2025). Briefly, upon removal from −80 °C storage, tissue slides underwent fixation in 4% PFA/PBS for 1 h at room temperature, followed by a PBS rinse and mounting of HybriWell-FL chambers. Dehydration comprised 5-min incubations in 70%, 85%, and 100% ethanol, followed by 1.5–3 h in fresh 100% ethanol at 4 °C; subsequent washes in PBST (PBS with 0.5% Tween-20) rehydrated the tissue.

Overnight reverse transcription at 37 °C in a humidified chamber used 50 μM random N20 primer, 1× RevertAid RT buffer, 20 U μL□^1^ RevertAid H Minus M-MuLV reverse transcriptase, 500 μM dNTPs, 0.2 μg μL□^1^ BSA, and 1 U μL□^1^ RiboLock RNase inhibitor. After PBST rinsing, cDNA immobilization used BS(PEG)_9_ stock diluted 1:4 in PBST for 1 h at room temperature. Crosslinker quenching consisted of a rinse in 1 M Tris-HCl (pH 8.0), a 30-min incubation in fresh Tris-HCl, and two PBST washes.

The gene panel comprised up to 12 padlocks per transcript, each encoding gene identity through a 7-nt index. Padlock hybridization and ligation proceeded for 30 min at 37 °C and 45 min at 45 °C in 1× Ampligase buffer supplemented with 100 nM of each probe, 0.5 U μL□^1^ Ampligase, 0.4 U μL□^1^ RNase H, 1 U μL□^1^ RiboLock, an additional 50 mM KCl, and 20% formamide. Two PBST rinses and one wash in hybridization buffer (2× SSC, 10% formamide) preceded a 15-min incubation with 1 μM RCA primer at room temperature; unbound primer was removed by three 2-min hybridization-buffer washes and subsequent PBST rinses.

Sequential rolling-circle amplification began with a 2-h incubation at 40 °C using phi29XT polymerase (New England Biolabs); each 1-mL reaction combined 600 μL nuclease-free water, 200 μL 5× reaction buffer, 100 μL 10 mM dNTP mix, and 100 μL polymerase. Direct replacement with a second mixture: 1× phi29 buffer, 0.25 mM dNTPs, 125 μM aminoallyl-dUTP, 0.2 μg μL□^1^ BSA, 5% glycerol, and 1 U μL□^1^ phi29 polymerase—supported further amplification for >12 h at room temperature. Following PBST rinsing, rolonies underwent a 1-h room-temperature crosslinking incubation with BS(PEG)9 diluted 1:4 in PBST, a 30-min quench in 1 M Tris-HCl (pH 8.0), and final PBST washes.

### Seq-o-Matic hardware

Seq-o-Matic combines a motorized fluorescence microscope, a heated three-slide perfusion chamber mounted on the microscope stage, and a reagent-exchange fluidics system, all controlled by one Python application (**<u>Fig. 1</u>**). Samples remain on the microscope stage throughout a run, and reagents are delivered through perfusion ports, so samples are never removed or remounted between cycles. All components are commercially available (**<u>Table S2</u>**). A more detailed assembly guide is provided in **<u>Supp File 1</u>** and at https://github.com/BarseqLab/Seq_o_matic/blob/main/docs/setup_guide/Seq-o-matic_Setup_Guide.md.

#### Microscope

We used a Nikon Ti2-E microscope with a motorized XY stage and Z drive, controlled through Micro-Manager 2.0 (Edelstein et al., 2014) and Pycro-Manager (Pinkard et al., 2021). Fine focus used a Nikon objective piezo, driven by an analog output of 0–10 V that corresponds to 0–400 µm. We imaged samples through the 1-mm-thick slides with a Nikon CFI S Plan Fluor LWD 20XC/0.70 NA objective. Excitation came from a Lumencor Celesta light engine through an X-Light v3 spinning-disk unit, and images were recorded on a Photometrics Kinetix camera with 3,200 × 3,200 pixels of 6.5 µm (0.33 µm per pixel at the sample with the 20× objective). The four sequencing channels used 514-nm excitation with 565/24-nm emission (G), 561-nm excitation with multiband emission (T), 640-nm excitation with 676/29-nm emission (A), and 640-nm excitation with 775/140-nm emission (C). Hybridization images were acquired at 488, 514, 561, and 640 nm excitation, with DAPI at 365 nm and a transmitted-light channel.

#### Imaging and heating chamber

Samples were held in a Bioptechs Tri-Imaging Chamber, a heated parallel-plate perfusion flow cell that holds three 25 mm × 75 mm slides in one stage adapter. From bottom to top, each chamber stack consisted of the stage insert, the tissue slide (tissue side up), a 0.25 mm thick inner silicone perfusion gasket that defines the flow channel over the tissue, a perforated indium tin oxide (ITO)-coated slide with its bus bars facing up, a 0.75-mm upper gasket, and the heated chamber top, which carries a thermistor on the inflow side. Reagents enter and leave through stainless-steel perfusion tubes (0.5-mm inner diameter for inflow and 1.0 mm for outflow). Current through the ITO layer heats the chamber. Each chamber was regulated by its own Bioptechs Delta T controller. A National Instruments DAQ supplied one analog setpoint shared by all controllers, read each controller’s temperature output, and switched each controller between heating and imaging modes, so that heating was off except during heated incubations.

#### Fluidics

Reagent bottles fed two cascaded 12-port selector valves (Elveflow MUX Distributor) through two 6-channel inline degassers (Biotech Fluidics Degasi Plus Classic). The first selector carried 11 reagent ports and a pass-through to the second, so that up to 16 reagents were addressable. The common outlet was split to three channels of a multichannel peristaltic pump (Ismatec ISM930C; PharMed BPT tubing, 0.51-mm inner diameter), which ran at 1.5 mL min□^1^per channel and delivered each reagent to the three chambers in parallel. The outflow went to waste. We have also tested a configuration in which a syringe pump with an integrated multiport valve (Tecan Cavro Centris with a 12.5 mL syringe) replaces the peristaltic pump. This configuration required less frequent maintenance in our hands but delivered reagents more slowly, because reagent has to be dispensed to the three chambers sequentially. All data in this study were acquired with the peristaltic pump.

### Seq-o-Matic software

#### Architecture

The control software (https://github.com/BarseqLab/Seq_o_matic.git) is a Python desktop application with a graphical user interface (GUI), a fluidics module, an imaging module, and device drivers for the pump, selectors, heaters, and microscope. Hardware and imaging settings are stored in JSON configuration files: serial ports, pump tubing diameter, the reagent-to-port map, heater DAQ lines, the exposure of each channel, pixel size, image size, tile overlap, focus and alignment channels, and stage directions. Laser intensities and filter settings are stored in the Micro-Manager configuration file. None of these settings can be edited from the GUI. Some of the device drivers were adapted from ACQ4 (Campagnola et al., 2014).

#### Protocol file

Each experiment was defined by a protocol file: an ordered list of fluidics cycles (e.g., primer addition, the first gene sequencing cycle, later gene sequencing cycles, hybridization, and stripping) and imaging cycles (reference, sequencing, or hybridization imaging). Users built the protocol file in a recipe-builder dialog. The software inserted a reference imaging cycle before the first imaging cycle if the user had left it out, and then executed the entries in order. Fluidics entries went to the fluidics module and imaging entries to the imaging module, and each entry started after the previous one finished. Before each run, the software checked every device and primed each reagent line with 1 mL of reagent, followed by 4 mL of PBST.

#### Fluidics module

Each fluidics cycle was defined by a recipe file, a list of steps of three types: pumping a set volume from a reagent port at 1.5 mL min□^1^, a heated incubation for a set time, and a room-temperature incubation. Consecutive steps were separated by 3 s, and the software waited 2 s for the selector to settle before each pumping step. Each recipe file was copied into the experiment folder when first used, so that every run kept a record of the recipes it executed. For heated incubations, the software switched the controllers to heating mode, polled each chamber’s temperature every 5 s until it reached the setpoint of 57.5 °C, held the setpoint for the specified time while logging the temperature every 10 s, and then returned the controllers to imaging mode. If a chamber did not reach the setpoint within 300 s, the software logged an error and stopped the run. When the devices were configured, the software also tested each heater with a 15-s heating pulse.

#### Imaging module

For each tissue section, the user defined four anchor positions at its corners during the reference imaging cycle. In each imaging cycle, the module first refocused at each anchor. It acquired a z-stack in the DAPI channel from −40 to +40 µm around the stored focus in 1.5-µm steps, scored the sharpness of each plane on the central 1,600 × 1,600 pixels (the mean gradient magnitude after a 3 × 3 median filter and a 3 × 3 sharpening kernel), and took the sharpest plane as the new focus. Focus changes of 20 µm or more were flagged as errors. The module then registered the best-focus DAPI image at each anchor to that anchor in the reference cycle by phase correlation, with a Blackman–Harris window, zero padding, and sub-pixel refinement of the correlation peak by fitting a quadratic surface to its 3 × 3 neighborhood. The median offset across the anchors of a slide was applied to every position on that slide. The module then tiled each section with image tiles overlapping by 10%, spanning the bounding box of the corrected anchors, and set the focus of each tile from a plane fitted to the four anchor focus positions. Each tile was imaged as a z-stack from −15 to +15 µm in 1.5-µm steps (20 planes). Sequencing cycles were imaged in the G (60 ms), T (30 ms), A (20 ms), and C (40 ms) channels, and the first cycle of each sequencing type also in DAPI (20 ms). Hybridization cycles were imaged at 488 (100 ms), 514 (30 ms), 561 (30 ms), and 640 nm (20 ms), in DAPI (20 ms), and with one transmitted-light plane. As each tile finished, a separate thread computed the maximum-intensity projection of each channel and saved it to network storage; the raw stacks were then deleted. The projections, recipes, logs, and position files were copied to a local image server. Each experiment folder also held a time-stamped operator log, the temperature record of each heater, the corrected positions of every cycle, and a table of the reagent volumes required for the run.

### Automated sequencing protocol

A gene-sequencing experiment consisted of sequencing primer addition, seven gene sequencing cycles with imaging after each cycle, stripping, and hybridization with fluorescently labeled oligonucleotides followed by hybridization imaging. Volumes below are per chamber, and heated incubations used the setpoint above. A more detailed running protocol is provided in **<u>Supp File 1</u>** and at https://github.com/BarseqLab/Seq_o_matic/blob/main/docs/setup_guide/Seq-o-matic_Setup_Guide.md.

#### Gene sequencing cycles

Sequencing was done with the Illumina MiSeq nano v2 kits (Illumina). In the first cycle, we incubated the sample with incorporation buffer (PR2; 1.5 mL, 3 min heated) and washed with PBST (PBS with 2% tween-20, 3 mL). We then blocked with iodoacetamide (3.1 mg/mL, 1 mL, 3 min heated), washed with PBST (1 mL), and exchanged into incorporation buffer (3 mL). We incubated with incorporation mix (IMS; 1 mL, 3 min heated) and washed four times with PBST (4 mL, each followed by a 3-min heated incubation) and once more with 4 mL of PBST. We then stained with DAPI (2 mL, 10 min at room temperature), washed with PBST (2 mL), and exchanged into scan mix (SRE; 1.5 mL) for imaging. Each later cycle began with cleavage, consisting of incorporation buffer (2 mL), cleavage mix (CMS; 1 mL, 3 min heated), and incorporation buffer (3 mL), and then repeated the steps of the first cycle from the PBST wash onward, with a 1-mL instead of a 3-mL first wash. Barcode sequencing cycles used the same steps with each IMS incubation performed twice, each preceded by 3 mL of incorporation buffer, and in cycles 2 onward with two 1-min CMS incubations.

#### Stripping, hybridization, and primer addition

To strip, we applied three rounds of strip buffer (60% formamide, 2x SSC, 0.01% Tween-20; 4mL per round), each followed by a 5-min heated incubation. For hybridization after stripping, we washed with FISH wash (2x SSC, 10% formamide; 5 mL), incubated with the hybridization oligonucleotide mix (0.25 µM per oligo in FISH wash; 1 mL, 2 min heated and then 10 min at room temperature), washed with FISH wash (3 mL), stained with DAPI (0.01 mg/mL in PBST; 4 mL, 10 min), washed with PBST (4 mL, 10 min), and exchanged into SRE (1.5 mL). To add sequencing primers, we washed with FISH wash (5 mL), incubated with the gene or barcode primer mix (1 µM oligo in FISH wash; 1 mL, 10 min at room temperature), washed three times with FISH wash (3, 1, and 1 mL, each followed by 2 min at room temperature), and washed with PBST (2 mL); primer addition after stripping included an additional 2-min heated incubation and a single 3-mL FISH wash in place of the three washes.

### Image processing

We processed the maximum-projection images with our BARseq image-processing pipeline (X. Chen et al., 2025), which is available with the Seq-o-Matic software. We first denoised each sequencing channel with noise2void (Krull et al., 2019), using a pre-trained model for each channel. For each cycle, we then corrected the shifts between channels, subtracted the background by morphological opening with a rolling ball of radius 6 pixels, and corrected channel bleed-through with a calibrated bleed-through matrix. The channel shifts and the bleed-through matrix came from a calibration of the instrument (X. Chen et al., 2019). We registered every sequencing cycle to the first sequencing cycle by translation. To do so, we summed the four channels, divided the image into 256 × 256-pixel blocks, and kept the brightest quarter of the blocks. We upsampled each kept block fivefold and masked pixels brighter than 2,000 intensity units, so that somata and bright debris did not dominate the alignment. We removed low-frequency background by subtracting a morphological opening (disk radius 10 upsampled pixels), averaged the cross-correlations across blocks, and applied the resulting translation, resolved to 0.2 pixels. We registered the hybridization image to the first sequencing cycle in the same way. Finally, we stitched the tiles of each section with MIST (Chalfoun et al., 2017) on two-fold-downsampled first-cycle images and used the stitching transforms to place rolonies and cells in section coordinates.

### Rolony decoding

We decoded gene rolonies with bardensr (S. Chen et al., 2021) using a codebook of seven-cycle barcodes, which for the mouse panel contained 106 genes, 101 of which had padlock probes in this panel, and five unused barcodes. We trimmed 160 pixels from each tile border, normalized each channel in each cycle by the median, across control tiles, of the maximum intensity in the central 20% of the tile, and ran bardensr with a noise floor of 0.05. We set the spot-calling threshold of each dataset to control the false discovery rate (FDR), which we estimated from the unused barcodes as the number of rolonies per unused barcode divided by the number of rolonies per gene barcode. We scanned ten thresholds within ±0.1 of an initial value (the median of the per-tile maximum evidence for unused barcodes) and chose the lowest threshold at which the mean FDR across control tiles was below 5%. We detected the three hybridization markers (*Slc17a7*, *Gad1*, and *Slc30a3* in the mouse panel) as regional maxima exceeding per-channel prominence and background thresholds (30, 20, 50, and 20 intensity units in the four channels).

### Cell segmentation and quality control

We segmented cells with Cellpose v0.6.5 (Stringer et al., 2021) using the generalist cytoplasm model, a cell diameter of 50 pixels, a flow threshold of 0.6, and a cell-probability threshold of 0. The cytoplasmic input was the sum of the first three hybridization channels and the first sequencing cycle, smoothed with a Gaussian filter (σ = 5 pixels), and the nuclear input was the DAPI channel of the hybridization image. We dilated each mask by 3 pixels without merging neighboring cells and assigned each decoded and hybridization rolony to the cell whose dilated mask contained it. Because neighboring tiles overlap, a cell could be segmented twice; we therefore treated cells from different tiles of the same section whose centroids fell within the same 5-µm box as duplicates and kept the copy with the most counts. We retained cells with at least 20 counts and at least 5 detected genes. For the mouse brains, counts and genes were summed over all codebook entries, including the three hybridization markers and the five unused barcodes; the unused barcodes carried 0.14% of all counts. For the marmoset, we applied the same thresholds to the 60 panel genes.

### Sequencing quality metrics

We compared the three datasets sequenced on the same Seq-o-Matic instrument (NOD, WSB, and marmoset) with four C57BL/6J brains sequenced manually on the same microscope, before it was automated (X. Chen et al., 2025). All metrics used the first seven sequencing cycles of the processed and registered images, reading the four channel intensities at the pixel of each rolony. In each cycle, the called base of a rolony was its brightest channel. We excluded the two C57BL/6J brains that were used in the strain analysis, because they were sequenced on a second Seq-o-Matic system with a different microscope, so their raw intensities are not directly comparable. A subset of the imaged tiles from each brain were used for quality metrics analysis.

#### Signal retention

Because restricting the analysis to decoded rolonies would select for rolonies whose signal survived all seven cycles, we measured retention on all rolony signals in the first cycle, whether or not they were decoded. We identified these signals as local maxima with a prominence of at least 50 intensity units in any first-cycle channel and took the union of their positions across channels. To correct for differences in brightness among channels, we divided the called-base intensity in each cycle by a per-channel scale factor, the median first-cycle called-base intensity of all signals called in that channel in that brain, and applied the same factor in all cycles. The retention of a signal at cycle *c* was its normalized called-base intensity at cycle *c* divided by that at cycle 1. We took the median across signals within each tile and then the median across tiles, excluding tiles with fewer than 200 signals, which were almost entirely off-tissue, and tiles in which retention was undefined in any cycle.

#### Quality score

We quantified the channel specificity of each decoded rolony in each cycle as the intensity of its brightest channel divided by the L2 norm of its four channel intensities, which ranges from 0.5 when all four channels are equal to 1 when all signal is in one channel. We took the median across decoded rolonies within each tile and then the median across tiles, excluding tiles without decoded rolonies or with an undefined score in any cycle.

#### Phasing

If each molecule in a rolony fails to extend with probability *p* per cycle, then after *n* cycles the fraction of molecules exactly one base behind is *np*(1 − *p*)^(*n* − 1), and the in-phase fraction at cycle *n* − 1 is (1 − *p*)^(*n* − 1). Their ratio, *R*(*n*) = *np*, grows linearly with *n*, and its slope estimates *p*; a fixed defect or a brightness-scaled background would instead give a residual that does not change with *n*. To estimate *R*(*n*), we used decoded rolonies whose base call changed between cycles *n* − 1 and *n* (*n* = 2,…, 7). To avoid confounding lagging signal with bleed-through, we used only the eight base changes between the two bleed-through groups (G→A, G→C, T→A, T→C, A→G, A→T, C→G, and C→T). We further required a quality score of at least 0.90 and a called intensity of at least 20 in cycle *n* − 1, and a distance of at least 5 pixels from any other decoded rolony. The residual was the intensity at cycle *n* in the channel called at cycle *n* − 1. Because the processed images contain no negative values, the background in an unoccupied channel is positive and scales with local brightness. We therefore subtracted from each residual a brightness-matched background from the same rolony: its cycle-*n* intensity in the bleed-through partner of the previous channel, scaled by the ratio of the two channels’ null levels in that tile. The null level of a channel was its mean intensity among rolonies that could not carry signal in it, estimated when at least 50 such rolonies were present. If the null level of the bleed-through partner could not be estimated, we subtracted its intensity without scaling; if that of the previous channel could not be estimated, *R*(*n*) was undefined for that tile. In each tile, *R*(*n*) was the summed corrected residual divided by the summed called intensity at cycle *n* − 1. We took the median at each *n* across the tiles in which *R*(*n*) was defined at all six transitions and fitted *R*(*n*) against *n* by least-squares regression with an intercept. For 95% confidence intervals, we used a hierarchical bootstrap (2,000 iterations). In each iteration we first resampled the eight base changes with replacement, pooling each tile’s residuals and denominators over the resampled base changes (a base change with fewer than 100 qualifying rolonies in a tile was not counted for that tile). We then resampled tiles with replacement among those with a defined *R*(*n*) at all six transitions, and refitted the line; iterations with fewer than 10 valid tiles were discarded. Resampling base changes treats them as independent estimates of the same quantity, so that their disagreement enters the interval. We report the median bootstrap slope with its 2.5th–97.5th percentile interval. The fraction of signal lagging by one base after seven cycles is *R*(7) = 7*p*. Per-brain confidence intervals for retention and quality score (Fig. 2C, D) came from 2,000 bootstrap resamples of tiles.

### Marmoset cell typing

We clustered the marmoset cells hierarchically in Scanpy (Wolf et al., 2018) using the 60 panel genes. We normalized each cell to a total of 10 counts and applied a log2(1 + *x*) transform. Before clustering, we replaced the *SLC17A6* count of each cell with its count minus its *GAD1* count, clipped at zero, to remove the effect of incomplete removal of the *GAD1* detection probes. For the first level (H1), we computed 50 principal components (PCs), built a 15-nearest-neighbor graph on the first 30 PCs, and ran Leiden clustering (Traag et al., 2019) at resolution 0.7, and we annotated the clusters as glutamatergic, GABAergic, or other types. For the second level (H2), we re-clustered the glutamatergic cells using 30 recomputed PCs, a 20-nearest-neighbor graph, and Leiden clustering at resolution 1.2. We did not apply batch correction. The spatial segmentation below used gene counts, not cell-type labels; cell types served only to identify where the bins leave the cortex, for which we counted a cell as non-cortical if its H2 type was a non-cortical glutamatergic type or its H1 type was the thalamic GABAergic type.

### Cortical binning and spatial domain segmentation

#### Section and bins

We analyzed one representative hemi-coronal section. Along the cortical ribbon, we manually placed 71 pairs of points, one on the pial surface and one on the white-matter boundary. Bin *i* was the quadrilateral whose corners were pairs *i* and *i* + 1, so that the 70 bins tiled the ribbon as an ordered chain of full-thickness columns (median tangential width 388 µm; range, 142–728 µm). We assigned each cell to the bin that contained its centroid; cells outside the ribbon were not binned.

#### Features

For each bin, we summed the counts over all cells in the bin that passed QC (20 reads per cell and 5 genes per cell). We smoothed these counts along the chain with a three-bin kernel that weighted the bin by 1/2 and each neighbor by 1/4, reflected at the ends of the chain. We converted the smoothed counts to proportions, applied the Hellinger (square-root) transform (Legendre & Gallagher, 2001), centered each feature, and reduced the profiles to 10 PCs.

#### Segmentation

We divided the ordered chain into *k* contiguous domains by exact one-dimensional dynamic programming (Fisher, 1958), in which the cost of a domain was its within-domain sum of squared deviations from the domain mean in the 10-PC space. For each *k* from 1 to 15, the dynamic program found the globally optimal borders and their total cost *W*(*k*).

#### Number of domains

Because *W*(*k*) always decreases with *k*, we compared the decrease gained by each additional domain with that expected if the domains of the current fit contained no further structure. For each step from *m* to *m* + 1 domains (*m* = 1,…, 14), we started from the optimal *m*-domain fit, pooled the cells of each domain, and refilled every bin with its own number of cells drawn with replacement from its domain’s pool. We then recomputed the features and the dynamic program. This null holds fixed the number of bins, the number of cells in each bin, the *m*-domain borders, and the processing, and randomizes which of a domain’s cells fall into which bin. We resampled cells rather than individual counts because counts from the same cell are not independent. We computed 500 surrogates per step. The excess gain of a step was the observed decrease *W*(*m*) − *W*(*m* + 1) minus the median surrogate decrease, set to zero when negative, and we plotted the cumulative excess up to each *k* as a fraction of the total excess up to *k* = 15 (Fig. 3C). We selected *k* = 5 at the shoulder of this curve, without a numerical threshold. As a separate test, we compared each observed decrease with the 95th percentile of its surrogates with Holm correction across the 14 steps; this test sets only a lower bound on *k*. We ran the analysis on the full chain of 70 bins (Fig. 3B, C) and on its cortical part alone (bins 1–63). The domains and cortical areas were defined by visual comparison of the sections to the marmoset atlas (Paxinos, 2012).

### Cell typing of the mouse strain dataset

We clustered cells from the four brains sequenced with Seq-o-Matic (two C57BL/6J, one NOD, and one WSB) together at three levels, following our previous hierarchical approach (X. Chen et al., 2025). All four datasets used the same codebook. **First level (H1).** We normalized each cell to 10 counts (CP10) and applied a log2(1 + *x*) transform, computed 50 PCs from all codebook entries, and integrated the brains with Harmony (Korsunsky et al., 2019) (batch variable, brain; θ = 2; at most 20 iterations). We built a 15-nearest-neighbor graph on the first 30 Harmony-corrected PCs and ran Leiden clustering at resolution 1.0. We assigned H1 clusters to glutamatergic, GABAergic, or other cells based on *Slc17a7* and *Gad1* expression and subclass marker genes. **Second level (H2).** We re-clustered the glutamatergic cells using 30 recomputed PCs, Harmony (θ = 2), a 20-nearest-neighbor graph, and Leiden clustering at resolution 0.6. We annotated H2 clusters manually based on marker genes and laminar position and named them after the corresponding subclasses in the Allen Brain Cell (ABC) atlas (Yao et al., 2023); two clusters with the same identity (both L4/5 IT) received a single label. **Third level (H3).** We re-clustered each H2 type separately with its own PCA (the smaller of 20 PCs and one-third of the number of cells), Harmony (θ = 1), a 15-nearest-neighbor graph, and Leiden clustering at resolution 1.0. All clustering used Scanpy with a fixed random seed. We used Harmony-corrected PCs only to define clusters; all other analyses used expression values without Harmony correction.

#### Quality control of H3 types

We excluded H3 types that were likely to reflect technical artifacts rather than cell types: types with few cells, low read counts, or spatial distributions inconsistent with a cell type. We evaluated each H3 type on five criteria: (1) fewer than 100 cells; (2) mean *Slc17a7* expression below 25% of that of its parent H2 type; (3) median counts per cell below 70% of that of its parent H2 type; (4) spatial concentration, that is, occupying fewer than half of the sections expected for its size (based on 2,000 random draws from the distribution of its parent H2 type across sections and tiles) or having more than 30% of its cells in one section at more than five times the expected fraction; and (5) over-representation of a single brain by more than 0.20 relative to a size-matched null (Holm-corrected p < 0.05). We classified an H3 type as low quality if it failed criterion 2, failed at least four criteria, or failed both criteria 1 and 3, and as small if it had fewer than 100 cells but normal read depth. We then inspected the spatial distributions of the flagged types and also classified those dominated by artifacts as low quality. Of the 101 cortical H3 types, 16 were classified as low quality or small; all comparisons across strains used the remaining 85.

#### Spatial plots, proportions, and embeddings

We plotted cells at their centroid positions, colored by H1 type (**<u>Fig. 4A</u>**) or by cortical H2 type (**<u>Fig. 5B</u>**); for **<u>Fig. 5B</u>**, we rotated or mirrored each section so that dorsal was up and the midline was on the right, and all sections were drawn at the same scale. For proportions (**<u>Fig. 5A</u>**), we calculated the fraction of each brain’s cells in each of the 10 cortical H2 types, renormalized within each brain over those types, and likewise for the quality-controlled cortical H3 types. We summarized differences across brains as the ratio of the largest to the smallest fraction among the four brains. For each cortical H2 type, we computed a UMAP embedding (McInnes et al., 2018) (minimum distance 0.01, spread 1.0) from PCs recomputed without Harmony and a 15-nearest-neighbor graph, and plotted 40,000 randomly sampled cells from quality-controlled H3 types in random order (**<u>Fig. 5C</u>**).

### Comparison with manually sequenced data

To compare the H3 types with previously identified cortical cell types, we transferred labels from the manually sequenced adult C57BL/6J brain (X. Chen et al., 2025) (465,872 cortical excitatory neurons annotated with 71 H3 types) to all excitatory neurons of the two automated C57BL/6J brains by k-nearest-neighbor (kNN) mapping. We normalized both datasets to CP10 and log2-transformed them. To correct for gene-specific differences in detection efficiency, we multiplied each gene in the reference by the ratio of its mean raw count in the query to that in the reference. We then z-scored each cell across genes and scaled it to unit length, so that Euclidean distance ranked reference cells by their Pearson correlation with the query cell. We found the 15 nearest reference neurons of each query neuron by approximate nearest-neighbor search as implemented in BiocNeighbors (AnnoyParam) and assigned the most frequent reference label. For each pair of de novo H3 type *i* and transferred type *j*, we computed the Jaccard index *n*□□/(*n*□+ *n*□− *n*□□) (**<u>Fig. 4B</u>**). We counted an H3 type as concordant if its best-matching manual type (the largest Jaccard index), among manual types reaching a Jaccard index of at least 0.05 with at least one cortical H3 type, belonged to the manual H2 type corresponding to its own H2 type. To check the correspondence of the H2 types with ABC atlas subclasses, we mapped the same queries in the same way onto the isocortical glutamatergic cells of the ABC atlas 10x Genomics v3 dataset (262,012 cells) (Yao et al., 2023) at the cluster level.

### Brain-to-brain neighbor analysis

To compare the technical variation between separately sequenced brains with the differences between strains, we measured, within each cortical H2 type, how often the nearest neighbors of each neuron came from its own brain and from each of the other three brains sequenced with Seq-o-Matic. *Neurons.* We used neurons with at least 40 counts across all 114 codebook entries; this excluded 11%–22% of neurons, most of them from the WSB brain. Within each H2 type and separately for each H3 type, we sampled equal numbers of neurons from the four brains (the number in the brain with the fewest neurons of that type; H3 types with fewer than 20 neurons in any brain were excluded), scaled the samples down proportionally to at most 5,000 neurons per brain, and pooled them. *Normalization and embedding.* We normalized raw counts to CP10 and log2-transformed them, without per-brain correction or Harmony, and computed 20 PCs on the pooled neurons. *Neighbor fractions.* For each neuron, we found its 40 nearest neighbors among all other pooled neurons by exact Euclidean search in PC space, excluding only the neuron itself. Entry (a, b) of the brain-by-brain matrix was the fraction of the neighbors of brain-a neurons that came from brain b; because the four brains were balanced, chance was 0.25. We averaged five independent balanced samples per H2 type and report means across the 10 H2 types. *Mixing index and test.* For each pair of brains a and b, the mixing index (M_ab + M_ba)/(M_aa + M_bb) is 1 when the two brains are interchangeable and 0 when they are fully separate. We tested whether the two C57BL/6J brains mixed more than the NOD and WSB brains with a one-sided exact sign test across the 10 H2 types, and estimated 95% confidence intervals from 10,000 bootstrap resamples of H2 types.

### Gene-level comparison of strain and cell-type differences

To compare the magnitude of strain differences with that of differences among H3 types, we measured how well each gene distinguished H3 types and how well it distinguished strains within each cortical H2 type, using only quality-controlled H3 types. *Normalization.* We normalized raw counts to CP10 and log2-transformed them, without depth equalization or per-brain correction, as in the neighbor analysis. *Stratified AUROC.* For each codebook entry other than the three hybridization marker genes (*Slc17a7*, *Gad1*, and *Slc30a3*), we computed a one-versus-rest AUROC for each level of one factor, comparing cells only within levels of the other factor. For H3 types, we compared each H3 type with the other H3 types within each strain. For strains, we compared each strain with the other two within each H3 type, pooling the two C57BL/6J brains. The AUROC was the Mann–Whitney *U* statistic summed over strata divided by the summed products of positive and negative cell numbers; we skipped strata with fewer than 40 cells and levels with fewer than 20 positive or 20 negative cells within a stratum. We summarized each gene by a direction-free score, 0.5 + max|AUROC − 0.5| over levels, separately for H3 types and for strains (**<u>Fig. 5E</u>**). We excluded the three hybridization marker genes because they were detected with a simple intensity threshold in the hybridization cycle, unlike the sequenced genes, which were decoded by bardensr with a threshold calibrated to a false discovery rate of 5% in each dataset; their counts therefore often differ substantially between samples for technical reasons.

### Statistics

Confidence intervals of phasing slopes came from the hierarchical bootstrap described above. The choice of the number of marmoset domains used a resampling null that re-ran the full feature and segmentation pipeline in every surrogate, with Holm correction across steps. The brain-to-brain neighbor analysis used a one-sided exact sign test across the 10 H2 types. We state the numbers of permutations, bootstrap iterations, and cells in each subsection above.

## Data and code availability

The Seq-o-Matic control software is available at https://github.com/BarseqLab/Seq_o_matic.git. The assembly protocol and tutorial are available at https://github.com/BarseqLab/Seq_o_matic/blob/main/docs/setup_guide/Seq-o-matic_Setup_Guide.md. The image-processing pipeline is available at https://github.com/BarseqLab/BARseq_processing.git. Analysis scripts and data are deposited at Synapse https://dx.doi.org/10.7303/syn78096899.

## Acknowledgements

We thank Anthony Zador, Michael Dworjan, Alex Vaughan, and Yan Li for discussions that inspired this work; Brian Long, Katelyn Kostello, Brannon Calabia, Dhruvi Joshi, and Militha Madur for discussions on implementation of the software. This work was supported by the National Institutes of Health (NIH) [ZIAMH002984 (J.A.B.), U01NS132161 (X.C., J.M.K, I.R.W, G.H., P.R.), DP2MH132940 (X.C.)]. A.Y.F and J.A.B are supported by the National Institute of Mental Health (NIMH) Intramural Research Program. A.Y.F is in part supported by the Commonwealth of Australia through an Australian Government Research Training Program Scholarship. The authors thank the Allen Institute founder, Paul G. Allen, for his vision, encouragement, and support. The NIH authors’ contributions were on data from the marmoset samples and did not overlap with the grant-funded portion of this study.

The contributions of the NIH author(s) are considered Works of the United States Government. The findings and conclusions presented in this paper are those of the author(s) and do not necessarily reflect the views of the NIH or the U.S. Department of Health and Human Services.

## Competing interests

The authors declare no competing interests.

## Author contributions

Conceptualization: X.C.; Investigation: A.Z., A.Y.F., A.P.A., M.B., C.T.D., S.D.H., A.H., T.E.O., A.A.O., X.C.; Methodology: A.Z., Z.M., J.L., S.C., A.G., C.Slaughterbeck, N.G., M.C.P.R., C.Shi, Y.-C.W., R.G., H.Z., J.M.K., A.C., X.C.; Resources: A.Y.F., M.M., J.A.B., G.H., I.R.W., P.R.; Validation: A.Z., A.Y.F., X.C.; Data curation: A.Z., A.Y.F., D.M.M., X.C.; Software: A.Z., A.Y.F., X.C.; Formal analysis: A.Z., X.C.; Visualization: A.Z., X.C.; Writing – original draft: A.Z., X.C.; Writing – review and editing: A.Z., J.L., A.Y.F., X.C.; Supervision: J.A., S.D.H., J.W., H.Z., J.A.B., G.H., J.M.K., I.R.W., P.R., A.C., X.C.; Project administration: G.H., J.M.K., I.R.W., P.R., A.C., X.C.; Funding acquisition: J.A.B., G.H., J.M.K., I.R.W., P.R., X.C.

## Supplementary Tables

**Table S1.** Hands-on time and total time for different experiments using manual sequencing or automated sequencing.

| Experiment Type | Cycle type numbers | Manual Sequencing |  | Automated, Seq-o-Matic |  |
| --- | --- | --- | --- | --- | --- |
|  |  | hands-on time | Total time | hands-on time | total time |
| Genes only (X. Chen et al., 2025) | Gene Sequence round x7, Hybridization round x1 | 12 hrs | 4 days | 2 hrs | 1.5 days |
| Genes only, two panels | Gene Sequence round x14, Hybridization round x4 | 27 hrs | 9 days | 2 hrs | 3.5 days |
| Genes and Projections (Sun et al., 2021) | Gene Sequence round x7, Hybridization round x1, Barcode Sequence round x15 | 35 hrs | 12 days | 2 hrs | 4.5 days |

*Table S2. Hardware and consumable list*

See TableS2.xlsx

*Table S3. Gene panel and Oligo list*

See TableS3.xlsx

## Supplementary Files

*Supplementary File 1. Installation guide and run protocol*

See Seq-o-matic_Setup_Guide.pdf

## Notes

https://github.com/BarseqLab/Seq_o_matic.git

### Competing Interest Statement

The authors have declared no competing interest.

