## Supplementary material for "Robust and accessible in situ sequencing with an open-source automated sequencer": Supp File 1

### Seq-o-Matic: Hardware Installation and Day-of Setup

*Installation and running guide to Seq-o-Matic. Guide written by Jacob Lapka.*

*Version 2026-10-18, for Seq-o-Matic 2.5. The latest version of this guide is at https://github.com/BarseqLab/Seq_o_matic/blob/main/docs/setup_guide/Seq-o-matic_Setup_Guide.md. The reagent calculator workbook it refers to, Automation_Workbook.xlsx, is in the same folder: https://github.com/BarseqLab/Seq_o_matic/blob/main/docs/setup_guide/Automation_Workbook.xlsx.*

---

#### Hardware Installation

**Reagents → Degassers → Selectors → 4-way splitter → Pump (3 lines) → Chambers/Samples → Waste**

Reagents flow in one direction, from the rack to waste (**Fig. 1**):

1. **Reagent tubes → degassers.** Each reagent tube in the fluidics rack feeds a dedicated 0.5 mm fluidics line (IDEX) that passes through the inline degassers (Biotech Fluidics) to remove dissolved gas before it reaches the sample (**Fig. 2A**, **B**).
2. **Degassers → selectors.** Degasser outlets connect to the selector valves, which choose which reagent is delivered at each step. To accommodate the extra reagents within our 12-inlet selector (Darwin Fluidics), the twelfth inlet is connected to the output line of the second selector (**Fig. 2B**). Reagents should feed into the proper channel of the selector as delineated in Seq-o-Matic. Refer to the repository documentation for more details about the specific connections.
3. **Selectors → 4-way splitter → pump.** Selector outputs join at a 4-way splitter, dividing flow into three parallel lines. The three lines run through the pump cassette (ensure pump flow is oriented in the correct direction). The fluidics tubing is connected to the flexible pump tubing (VWR, MFLX95714-18) using a conical adaptor (IDEX, p-794) with the conical end fitted snugly into the pump tubing (**Fig. 2C**).
4. **Pump → sample chamber tops.** The downstream pump tubing is connected back to 0.5 mm fluidics tubing which is then connected to the input fitting of each chamber top on the stage (**Fig. 2D**, **E**). When samples are not present, this input tubing can be connected directly to the 1 mm output tubing (IDEX).
5. **Chambers → waste.** Each chamber output line runs directly to its own waste bottle. Be aware that that some sequencing reagents are toxic and the waste should be treated as such.
6. **Heaters.** The heaters (Bioptechs) are connected to the chamber top only after all fluidics setup is finished.

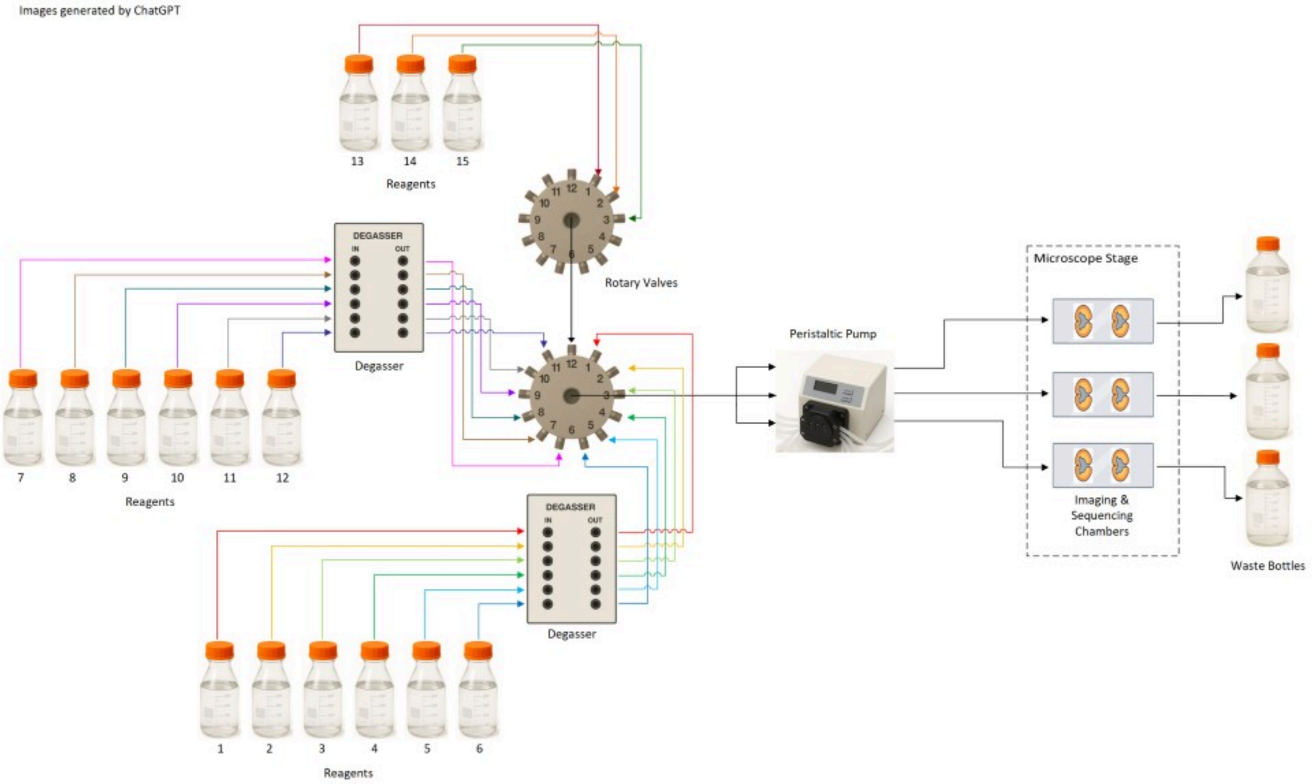


**Figure 1. Schematic demonstrating microfluidic flow throughout the perfusion system.**

#### Microfluidic Care

- **Fluidics tubing:** all tubing upstream of the sample is 0.5 mm (IDEX, 1516xl) while the output from the sample to waste is 1 mm (IDEX, 1517L). Whenever changing connections, avoid overly twisting the pump tubing as this can alter the flow rate or lead to leaks.
- **Reagents:** ensure that all reagent fluidic tubing correctly connects to the proper inputs, taking extra care at the degassers and selectors when inlets and outlets change.
- **Pump tubing:** do not twist or kink it. Ensure the orientation of the cassettes matches the direction of flow. Seat the cassettes on both sides until it clicks and unclip it after each run to prevent kinks. The most likely cause of uneven flow between chambers is distress to the pump tubing, either through over-twisting, extended use, or leaving clipped to the pump while not in use.
- **Exercise pump tubing:** anytime the pump tubing is changed out, exercise the lines to stretch them out until they flow consistently. Run water through them for 30 minutes – 1 hour. This evens out the flow rate between different pump lines.
- **Ferrules:** use a ferrule (IDEX) on every junction with 1/16″ VacuTight™ fittings (Cole Parmer), oriented correctly (tapered end toward the fitting). A missing or improperly installed ferrule can result in leaks. Replace distressed ferrules (check that they are properly seated at the end of the tubing and that the line is not pinched) at active junctions each run.
- **Chamber connections:** input line to input side, output line to output side. Attach fittings onto the metal pipe of chamber top prior to hooking up samples to fluidics (**Fig. 2D**). This provides a secure connection to add the input tubing without damaging the chamber top.

- **Liquid handling:** reagent tubing must reach the tube bottoms, with lids only lightly screwed on (tight lids pull a vacuum). Waste lines must not touch waste liquid, as this can result in backflow. Clean any drips immediately.
- **Software recognition:** check that the COM ports assigned for each device match the config files in Seq-o-Matic.
- **Wash tubing and reusable equipment after every run.** Seq-o-Matic is equipped with a "wash tubes" function that runs 4 mL (per chamber) of fluid through every line. It is recommended to wash with water in place of every reagent after every run to prevent particulate or chemical build-up within the tubing system.
- **Maintain microfluidic system:** the frequency of system maintenance varies with use and level of preventative care. For example, we have developed a schedule that calls for a 0.2% sodium azide wash (using the "prime lines" function) every two months after changing out all the tubing. Always follow these washes with two water washes using the "wash tubes" function. We change out all tubing, fastenings, and fittings every 3 – 5 months to prevent potential points of failure.
- **A comprehensive list** of the different consumables and components discussed in this guide can be found in **Table 1**.

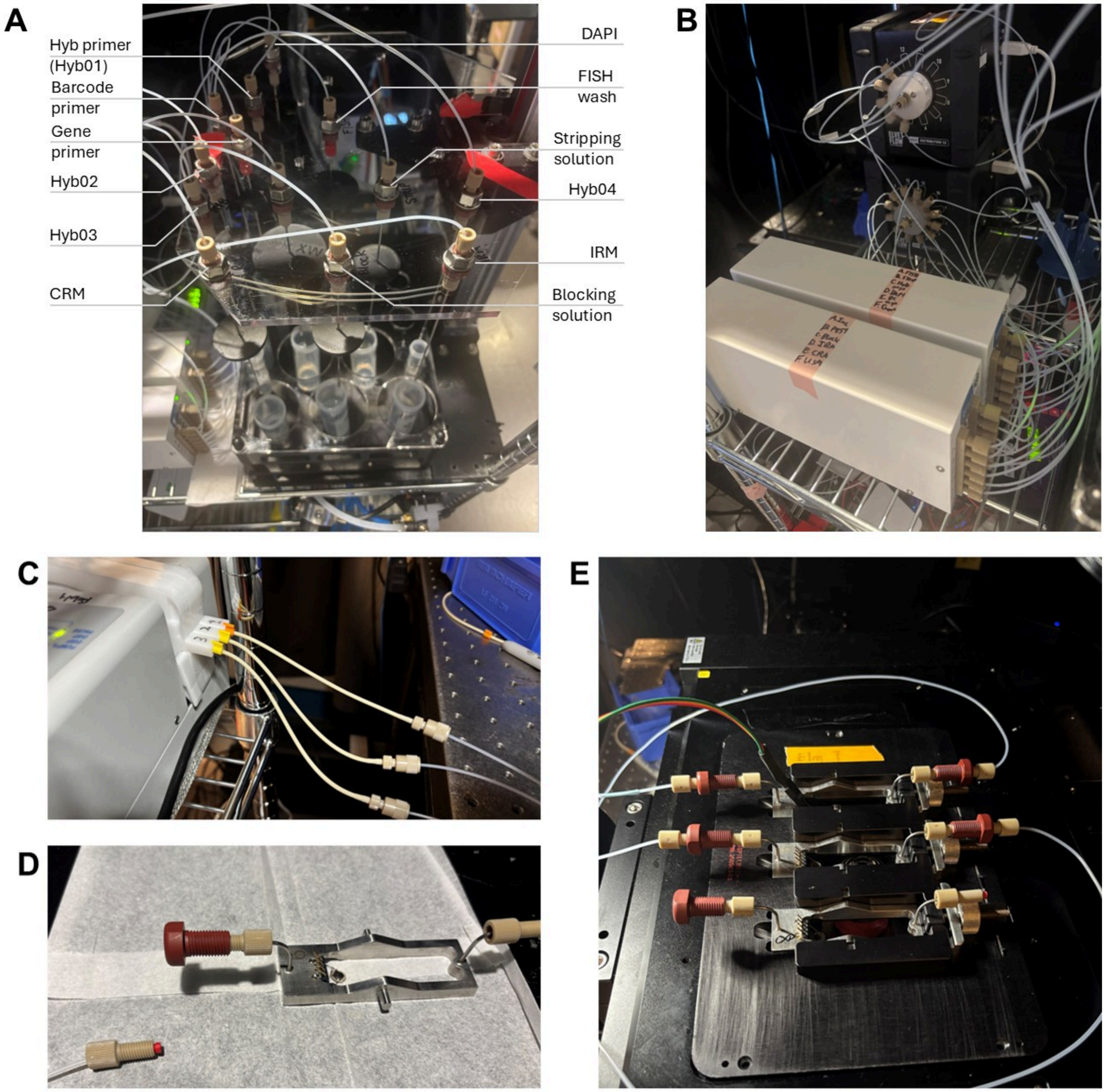


**Figure 2. Example images detailing most of the components necessary for Seq-o-Matic microfluidic setup.** **(A)** Reagent fluid rack. Each reagent has dedicated placement and tubing lines. **(B)** Reagents used most are connected to degassers (front). The Selectors (back) each contain 12 channels. Channel 12 of Selector 1 (back, bottom) connects to the middle chamber line of Selector 2 on top of it. The tubing for reagents that are infrequently used and in smaller volumes (such as additional hybridization primers) are hooked up directly to Selector 2 (channels 2, 4, and 5). **(C)** The output from the pump demonstrates the different flexible tubing and the orientation of the conical connection adaptor required. **(D)** Image of fitting assembly on custom chamber top. The left side of the chamber top, with the heating component (4

spikes) and thermistor (circular component inside the chamber space), is the input side, while the right is the output. The tan fitting is connected to the red Bulkhead Assembly with a ferrule (red) pointed towards the fitting. The output side has only a fitting and a ferrule. This setup allows easy insertion of samples into the fluidic system. **(E)** Three-slide stage insert with connections in 3 levels of attachment to fluidic system necessary for Seq-o-Matic run. (Top) Input and output tubing, along with the temperature control line attached above the thermistor. (Middle) Temperature control line not yet connected. (Bottom) Fluidics tubing not yet connected. While not connected to chambers, the input and output lines are connected with the same fitting-bulkhead union-fitting configuration used for chambers.

---

#### Sequencing Automation Setup and Operation

*For more detailed information regarding the Seq-o-Matic architecture and calibration, please refer to the repository README and docs/architecture.md.*

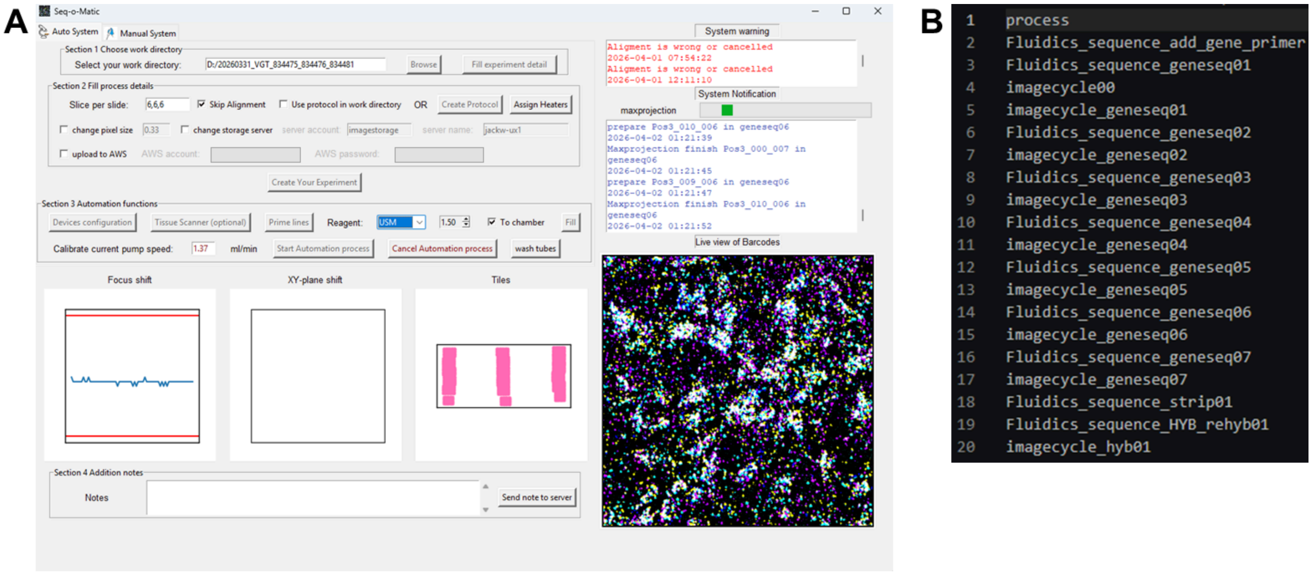


**Figure 3. Seq-o-Matic provides a GUI with easily adjustable settings that run customizable protocols. (A)** Example image of the Seq-o-Matic GUI during a sequencing run. **(B)** Example protocol.csv file consisting of 7 complete sequencing cycles and 1 hybridization cycle.

##### A. Software and data

1. **Restart** the control computer at least every 2 weeks; a full memory cache slows imaging.
2. **Start Micro-Manager** and load up the microscope configuration.
3. **Open Seq-o-Matic.** Use a terminal to open the `code/` directory of Seq-o-Matic and run `python mainwindow.py` to launch the application GUI (**Fig. 3A**).
4. **Create the experiment directory** on the first data drive and place the protocol (protocol.csv) in it. This file contains the automation sequence to follow based on primers, probes, and conditions used in the experiment. The format starts with "process" and must include "imagecycle00" prior to the

first round of imaging to properly mark the imaging field and set the focus. An example of 7 gene sequencing cycles followed by 1 hybridization cycle can be found in **Fig. 3B**.

5. **Load the correct reagent-sequence files.** Copy the default reagent.json files over the same-named files in the Seq-o-Matic reagent sequence folder. This ensures that every time, the correct .json files are using the correct files, and it can be time consuming to manually check the files every time. These files only change when altering the chamber size and amount of fluid exchange needed for the sample. Otherwise, they should remain as the default files.
6. **Configure the experiment** in Seq-o-Matic:
    - Select the directory and enter experiment details (to edit later, edit the .csv in the directory).
    - Enter slices per slide (e.g., 3,2,2). These refer to the number of image grids that will need to be created in micromanager and typically match how many samples are on each slide. Each slice will readout out as a position.
    - Check "Skip Alignment." The current system is stable enough that the xy alignment per cycle is not needed. However, some custom setups may not be stable enough and will instead want to utilize this option.
    - Check "Use protocol in work directory". This will prompt Seq-o-Matic to search inside the experiment directory and read the protocol.csv file. This is then the file that will be used during the sequencing run. Otherwise, you can create a protocol using the GUI.
    - If using less than 3 chambers, "Assign Heaters" allows for the selection individual temperature control units to run. If this is skipped while there are less than three heaters hooked up, Seq-o-Matic will read out that the heater is unable to raise the temperature and will run into an error.
    - Set the storage server. There are options to either upload the image output files to a local server, or alternatively, upload the files directly to AWS. For detailed information on how to input the specific pathways for these options, please refer to Setting up another lab (storage.json) in the README.
    - Click "Create Your Experiment," everything below it is disabled until you do. *If you later edit anything prior to this step (such as slices per slide or the protocol.csv file), click this button again and reconfigure devices.*
7. **Configure devices** from the device pull-downs. The heater must be plugged in to configure; if the stage chamber is not ready, configure it later (step 19).
    - If configuration fails, check that ports recognize the correct devices. Unplug and replug the USB hub, and restart Seq-o-Matic if needed.
8. **Confirm storage:** usually at least 5 TB free on the first data drive (for raw data) and 7 TB on the second drive (for max projection data) are effective buffers. The first must contain enough space for a full cycle of image stacks from which maximum projections are made and transferred to the second data drive. The second drive receives these image files for every cycle.

#### B. Reagents

9. **Prepare reagents** as described in the companion worksheet calculator (Automation_Workbook.xlsx):
    - Adjust the number of chambers and input the exact protocol for the run.
    - The number and type of cycles are used to calculate the total volumes necessary from set values per cycle type.

- For bulk reagents (such as PBST or FISH wash), filter the reagents prior to use to prevent small particles building up and clogging the fluidics tubing. Label every tube with name, volume, and date.

10. **Record starting reagent volumes** in a tracking sheet to refine future calculations and reduce waste.

#### C. Fluidics

11. **Load the reagents** into the fluidics rack and check that:
    - Each tube is in its correct position and holds at least its labeled volume.
    - Fluidics tubing touches the bottom of every tube (to prevent running dry).
    - Lids are lightly screwed on, and any drips are cleaned immediately.
12. **Empty waste bottles.** Keep waste lines above the liquid surface.
13. **Clip in the pump cassettes.**

> When spreading the fluidics across multiple lines, it is easy to hit the maximum flow rate of the pump. Seq-o-Matic compensates for this by varying the pumping time based on the measured flow rate.

14. **Calibrate current pump speed** to ensure even reagent flow across every line:
    - Weigh three empty receiving tubes.
    - In Seq-o-Matic, set the Reagent to PBST and volume to 1.5. Set the current pump rate to 1.5 mL/min. Press “Fill” to load the lines with PBST, pumping PBST through the lines for 1 min. It might take a couple times before it makes it through the microfluidic system into the waste.
    - Place the output tubing into each receiving tube and press “Fill.”
    - After the pumping is finished, weigh the now filled receiving tubes.
    - Subtract the tube weight and convert weight to volume to measure flow rate of each line. Record the average flow rate (mL/min). Each line should be within 10% of the mean for even flow.
    - If flow rate is too slow (below ~1.2 mL/min) or there is a major discrepancy between lines, check whether there is something clogging the tubing or is some other distress to the fluidic tubing. Consider changing the pump tubing out or recalibrating the pump directly to get to increase flow. Low flow rates slow the sequencing cycles.
    - Repeat these steps if any changes are made.
15. **Update the measured flow rate** recorded in the previous step in Seq-o-Matic under “Calibrate current pump speed.”
16. **Prime lines** (about 15 min; uses about 1 mL per line per chamber, followed by 4 mL of PBST). This ensures even perfusion across the sample as the selector switches between reagents.

#### D. Chambers and stage assembly

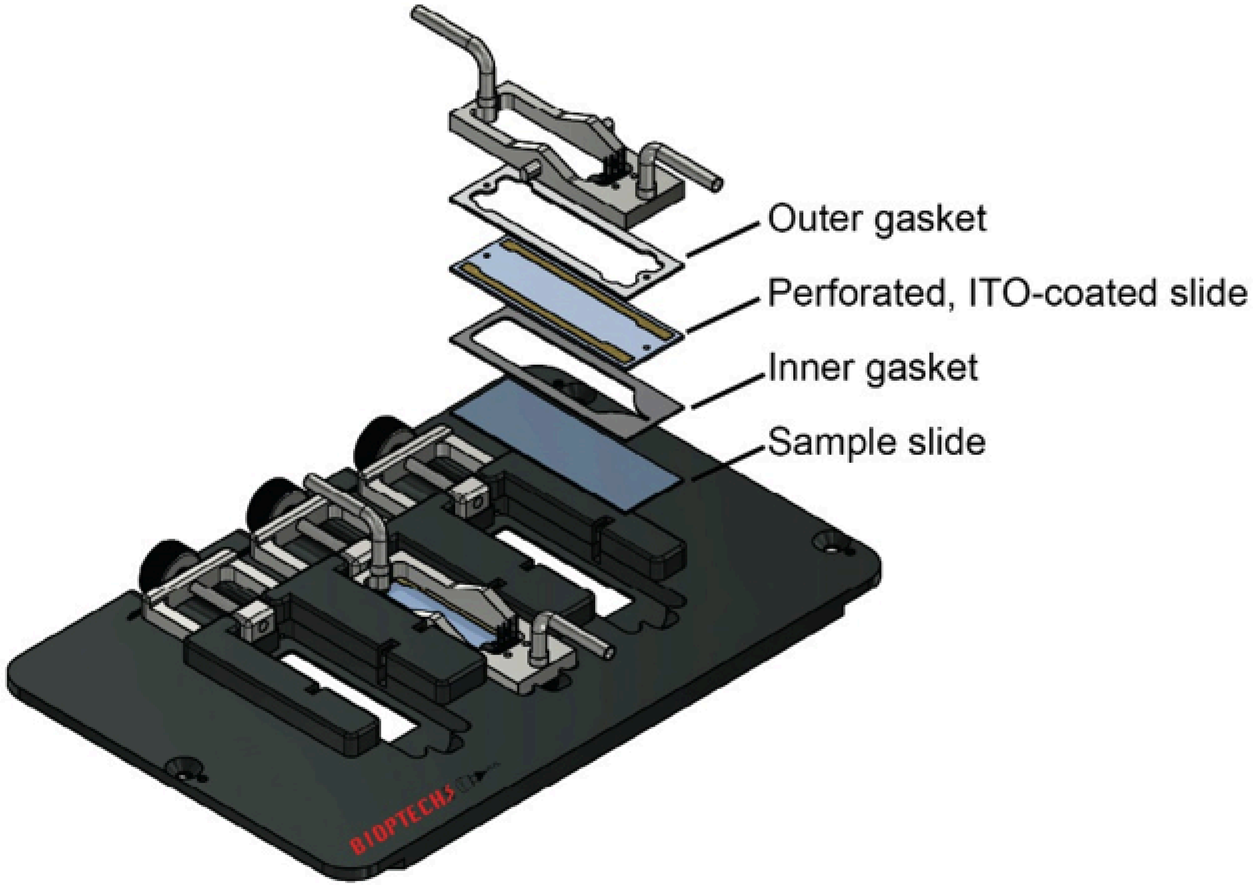


**Figure 4. Diagram of three-slide stage insert with chamber components shown.**

17. **Assemble each chamber** (disassemble the library-prep chamber first):
    - ITO-coated slide upside down (coating down) in the stage holder, long bus bar on the input side (away from the tightening mechanism).
    - Place a 0.25 mm perfusion gasket with the pointed side at the input.
    - Sample slide tissue-down inside the gasket, with the emptier side of the slide toward the input.
    - Flip the "sandwich" by holding the two slides together and carefully switching it so that the ITO slide is face up.
    - Dry the ITO slide with ethanol and wipe twice; add the 0.75 mm upper gasket (must be dry, replace if in doubt).
    - Place the chamber top with the thermistor on the input side and hand-tighten the screw as tight as possible.
    - As shown in **Fig. 2D**, first place a fitting on the pipe, followed by a red ferrule, and lastly add the bulkhead assembly (IDEX), pulling gently on both ends while tightening, so the pipe-to-chamber joint is not damaged.
    - Fill the chamber with DAPI by syringe (connected to tube with fitting and ferrule on end so it can screw into the chamber fittings).

> **Common failures:** cracked slides (clear debris), leaks (loose chamber or missing ferrule), an unflipped sandwich, any flipped component, and moisture in the upper gasket (blocks heating and degrades the ITO coating).

18. **Move stage insert** and chambers to microscope.
19. **Configure the heater before plumbing.** Plug in the heater cable firmly. A failed configuration may mean a faulty chamber top, which is easier to swap before lines are attached. Sometimes during configuration, the thermistor does not yet register a temperature change in the slide, and an error will display that the device failed to raise the temperature. If this is the case, configuring the heaters again will typically solve the issue. This is not an issue during runs as the heating step is much longer than a few seconds.
20. **Plumb the chamber:** replace screws and ferrules at active junctions, connect input and output lines (see Hardware Installation and **Fig. 2D, E**).
21. **Fill with PBST** (select the reagent, "Fill") and clear any bubbles in the lines and at the new connections:
    - Hold the chamber with the output up and tap until bubbles clear (tap against a sturdy shelf or with hand). Careful not to damage any of the lines or chambers.
    - Check for leaks and repeat until no bubbles remain.
22. **Secure the stage insert:**
    - Mount the stage insert onto the stage.
    - Move the stage to its extremes to confirm lines are slack and will not be pulled out; tape lines down clear of moving parts. Ensure the objective will not be damaged during the imaging cycle due to contact at extreme edges.
    - Confirm that the insert is level with the stage (not necessarily with the earth).
23. **Set imaging points:**
    - In Micro-Manager, open multi-dimensional acquisition, then "Edit positions".
    - Begin the live view, using the DAPI channel to get samples into focus.
    - For each ROI, mark the top, right, bottom, and left positions *in that order*. This will set the imaging grid for each slice.
    - Repeat this for each slice as marked in the “slices per slide” during Step 6.
    - Save as cycle00.pos in the experiment directory and confirm points are positioned correctly and in focus.
    - Point numbers must stay sequential and in order or imaging will fail (pos0-3 for first slice, pos4-7 for second, etc.).

#### E. Start and takedown

24. **Final checks:**
    - Protocol and sequence files correct.
    - If any information/step prior to pressing “Create Your Experiment” has changed, the button was pressed again and the devices reconfigured.
    - Flow rate is updated to match measured values.

- Waste containers are empty.
- Data storage on both drives is adequate.
- Chambers bubble- and leak-free and tightly screwed to the holder.
- Imaging cabinet lights are off.
- Both degassers are on with a steady green light.

25. **Click "Start Automation process"** and confirm the first step is correct.
    - Stay at least 15 min to catch potential leaks.
    - Stay until imaging of cycles 00 and 01 has started to check focus at the extremes; if any are off, cancel the run and fix the imaging points. Restart the run with the protocol beginning at imagecycle00.
26. **Check in** everyday while the sequencing run is ongoing. Check that the images are still in focus between cycles and that there are no leaks or other unexpected issues occurring.
27. **Takedown:** only after confirming that the run is finished and data are uploaded:
    - Connect each input line to its output line and swap reagent tubes for water tubes.
    - Press “Wash tubes” to flood each fluidic line with water (about 45 min; about 4 mL per reagent per chamber).
    - While the lines are washing, remove the sample chambers from the stage. The samples can be saved in PBS at 4°C in the event that they are still needed for referencing. Clean any reusable components thoroughly, as necessary based on the reagents used, and rinse them with water when finished.
    - Record final reagent volumes to refine calculations and reduce waste.
    - Close Micro-Manager and Seq-o-Matic.
    - Unclip the pump cassette to prevent kinks.
    - Empty waste containers.

---

#### Table 1

| Item | Supplier | Part Number |
|---|---|---|
| Peristaltic Pump 4 Channel | Masterflex | MFLX78018-20 |
| MUX Switch Distributor | Darwin Fluidics | LVF-MUX-D-12 |
| Pressure Sensor (optional) | IDEX | I2C PS200F |
| 6-Channel degasser (degasi plus classic) | Biotech fluidics | 00036356-A |
| ETFE Tubing 1/16" OD x .020" ID | idex | 1516xl |
| ETFE Tubing 1/16" OD x .040" ID | idex | 1517L |
| VacuTight™ Ferrule ETFE 1/16 Red | Idex | P-840 |
| VacuTight™ Fittings 1/4-28, 1/16" | Cole parmer | EW-02017-61 |
| Adapter Assembly, Conical | idex | p-794 |

| Item | Supplier | Part Number |
|---|---|---|
| Bulkhead Union Assembly | idex | p-441 |
| Cross Assembly (4-way splitter) | idex | p-722 |
| Masterflex® Ismatec® Pump Tubing, 3-Stop, PharMed® BPT, 0.51 mm ID | VWR | MFLX95714-18 |
| Delta T dish controller | Bioptechs | 0420-4-03 |
| Tri-imaging chamber base plate | Bioptechs | 200903-03 |
| Imaging chamber top | Bioptechs | 200903-1616 |
| Sample prep base | Bioptechs | 91603-0302 |
| Sample prep top | Bioptechs | 91603-1616 |
| non-coated microaqueduct slide for sample prep chamber | Bioptechs | 130119-5-200903-NC |
| ITO perfusion slides | Bioptechs | 130119-5-200903 |
| Silicone gaskets upper 0.75mm | Bioptechs | 1907-R22982B-750 |
| silicone gaskets perfusion 0.25mm | Bioptechs | 1907-R22982A-250 |
| silicone gaskets perfusion 0.5mm | Bioptechs | 1907-R22982A-500 |

**Table 1. Hardware and consumables list**

---

#### Helpful links

- Seq-o-Matic GitHub repository: https://github.com/BarseqLab/Seq_o_matic
- Reagent Calculator: Automation_Workbook.xlsx
